# Physiologic variation in sperm miRNAs tune embryonic gene regulatory programs and developmental outcomes

**DOI:** 10.64898/2026.03.04.709527

**Authors:** Grace S. Lee, James Garifallou, Samantha L. Higgins, Katherine Z. Scaturro, Adele Harman, Taylor Miller-Ensminger, Madeline N. Lamonica, Liana Savarirayan, Michael C. Golding, Colin C. Conine

**Affiliations:** Pharmacology Graduate Group, University of Pennsylvania Perelman School of Medicine, Philadelphia, PA, USA; Department of Genetics and Pediatrics, University of Pennsylvania Perelman School of Medicine, Philadelphia, PA, USA; Division of Neonatology, Children’s Hospital of Philadelphia, Philadelphia, PA, USA; Interdisciplinary Graduate Program in Genetics and Genomics, Texas A&M University, College Station, TX, United States; Department of Veterinary Physiology and Pharmacology, College of Veterinary Medicine and Biomedical Sciences, Texas A&M University, College Station, TX, USA; Transgenic Core, Children’s Hospital of Philadelphia, Philadelphia, PA, USA

**Keywords:** epigenetic inheritance, miRNA, sperm, embryonic development, alcohol

## Abstract

Small RNAs delivered by sperm can transmit environmentally regulated, epigenetically inherited phenotypes to offspring, yet the mechanisms by which modest changes in sperm microRNA abundance overcome dilution within the much larger egg to influence embryonic development remain unresolved. Here, we show that physiologically relevant variation in individual sperm miRNAs is sufficient to quantitatively program embryonic gene expression and developmental outcomes. Using parthenogenetic and fertilized embryos, we show that as few as 200 molecules of *miR-200c-3p* or *miR-465c-3p* induces reproducible, dose-dependent gene expression responses across defined developmental windows. Parthenogenetic embryos faithfully recapitulate early miRNA-driven gene expression changes observed in fertilized embryos, validating their use for isolating early regulatory mechanisms. We further developed AGO2-REMORA, an RNA adenosine base editor fused to Argonaute2 to map miRNA–mRNA interactions in embryos, revealing that early mRNA repression reflects direct miRNA targeting, while transcriptional changes at later stages arise as secondary consequences of these initial interactions. Furthermore, we show that modest elevation of *miR-200c-3p* during early development is sufficient to induce transcriptional alterations through early development and produce craniofacial phenotypes in late-stage embryos, recapitulating features of fetal alcohol syndrome associated with paternal alcohol consumption. Together, these findings establish a generalizable framework by which small perturbations in sperm miRNA content quantitatively modulate early gene regulatory programs, triggering cascades that persist throughout development and influence offspring phenotype.

## INTRODUCTION

Epigenetic information carried by sperm enables paternal environmental exposures to shape offspring phenotypes. Over the past two decades, studies in mammals have shown that paternal diet, stress, and toxicant exposure can yield both adaptive and maladaptive outcomes in progeny. Among the epigenetic factors implicated in this process, sperm microRNAs (miRNAs) have emerged as a key molecule for transmitting a father’s environmental experiences to his offspring^1^. Despite accumulating evidence that sperm miRNAs transmit paternal effects, the mechanistic function of these miRNAs during early embryogenesis to reprogram development and modulate inherited phenotypes remains unresolved. Fertilization creates a pronounced dilution problem because the sperm delivers a small payload into the vastly larger egg cytoplasm. This “stoichiometric barrier” raises the central mechanistic question of how the small input of sperm miRNAs can exert meaningful regulatory effects. Nonetheless, zygotic microinjection experiments have revealed that sperm miRNAs can transmit biologically meaningful signals that influence offspring phenotypes. For example, paternal stress in mice elevates specific sperm miRNAs, and microinjection of a cocktail of these miRNAs into one-cell zygotes recapitulates the offspring’s stress-related phenotypes^3,4^. Similarly, exercise^5^ and changes in diet^6^ also induce changes in the sperm miRNA payload, and when applied to naïve zygotes, are sufficient to recapitulate altered offspring outcomes. These findings establish that paternally delivered miRNAs function in the zygote, mediating post-fertilization gene regulatory events that alter developmental trajectories. Therefore, despite the dramatic difference in size and RNA content between sperm and egg, mounting evidence indicates that sperm miRNAs can influence early embryonic gene regulation and alter offspring phenotypes.

A well-characterized model of intergenerational epigenetic inheritance arises from preconception paternal alcohol exposure, which induces fetal alcohol syndrome-like craniofacial growth defects in mouse offspring^7^. In parallel, paternal alcohol consumption has been shown to remodel the sperm small RNA payload, including increased abundance of *miR-200c-3p* and decreased abundance of *miR-465c-3p* in sperm from alcohol-exposed males (Michael Golding Lab, *In revision*). Together, these observations establish a context in which defined, exposure-associated changes in individual sperm miRNAs can directly be tested for their capacity to reprogram early embryonic gene expression and influence developmental outcomes.

To date, much of the evidence for sperm miRNA function has been correlative, linking environmentally induced changes in sperm miRNA abundance with offspring phenotypes, and in some cases, to altered miRNA expression in somatic tissues of the progeny. One proposed mechanistic explanation for these findings is that specific sperm miRNA families, such as *miR-34/449,* may initiate a positive feedback loop, sustaining their own expression during embryogenesis and development^8^. However, this model does not address whether sperm miRNAs can act more broadly and directly in the early embryo to initiate gene regulatory changes.

Here, we propose a cascade model in which sperm miRNAs transiently act directly in the early embryo, downregulating targets through canonical base-pairing interactions. Repression of these early targets trigger indirect, downstream changes that alter developmental trajectories, thereby leading to persistent phenotypic outcomes. In this study, we tested this model by quantitatively elucidating how specific miRNAs influence early embryonic development. To do this, we microinjected defined quantities of *miR-200c-3p* and *miR-465c-3p* into zygotes and profiled gene expression at the late 2-cell, 4-cell, and morula stages in parthenogenetic and sperm-fertilized embryos to determine how physiologic variation in sperm miRNAs quantitatively alter embryonic gene expression programs. To distinguish direct miRNA targeting from downstream regulatory effects, we adapted REMORA (RNA-encoded molecular record of RNA-protein interactions) by coupling it to Argonaute2 for use in preimplantation embryos, enabling identification of miRNA target engagement in 2-cell parthenogenetic embryos to define immediate molecular responses and direct targets of sperm miRNAs following fertilization. Finally, we demonstrate that delivery of as few as 200 molecules, a dose within the endogenous range delivered by sperm, of a single sperm miRNA, *miR-200c-3p*, is sufficient to induce craniofacial defects in offspring, establishing direct causality between sperm miRNA perturbation and alcohol-associated developmental phenotypes.

## MATERIALS AND METHODS

### Animals

Male (8-12 weeks old) and female (5-7 weeks old) FVB/NJ mice were derived from breeders obtained from Jackson Laboratory (Strain #001800) and maintained in-house. All animal procedures were in strict accordance with the guidelines of the Children’s Hospital of Philadelphia and the University of Pennsylvania Institutional Animal Care and Use Committee regulations (CHOP IACUC Protocol #23-001364, UPenn PSOM IACUC Protocol #806911).

### Parthenogenesis and Embryogenesis

Mouse parthenogenesis and embryogenesis were performed as previously described^9^. Briefly, eggs were isolated from female mice superovulated using pregnant mare serum gonadotropin (PMSG) and human chorionic gonadotropin (hCG). For parthenogenesis, eggs were dissociated from cumulus cells using Hyaluronidase (3 mg/mL) and activated using KSOM with 10 mM SrCl_2_, 4 mM EGTA, and 0.05 μg/mL cytochalasin B for 1 hour. In vitro fertilization (IVF) was performed in HTF with 1 mM glutathione with capacitated sperm for 3 hours. Embryos were cultured in KSOM under standard conditions 37°C, 5% CO_2_, and 5% O_2_ to the indicated developmental stages. Detailed step-by-step procedures are available in Lee et al., *STAR Protocols*.

### Caudal sperm isolation for small RNA sequencing

Small RNA sequencing from sperm was performed largely as previously described^10^. Briefly, total sperm RNA was isolated and size-selected for small RNAs. In contrast to the original protocol (18-40 nucleotides), libraries in this study were size selected for 18-24 nucleotide fragments to enrich specifically for mature miRNAs. Libraries were prepared using a modified Illumina Tru-Seq small RNA library prep protocol and sequenced on an Illumina NextSeq 1000/2000 platform. Libraries were mapped and normalized to reads per million as described previously^10^. Raw read count data and normalized data are provided (Supplemental Table 1).

### Synthetic miRNAs

Chemically synthesized small RNAs corresponding to the mature guide strand sequence of *miR-200c-3p* or *miR-465c-3p* and the complementary passenger strands were obtained as single-stranded oligonucleotides (IDT). The guide strand contained a 5′ monophosphate to promote preferential loading into Argonaute, while the passenger strand was synthesized without a 5′ phosphate. A single–nucleotide mismatch was introduced into the passenger strand to further bias strand selection toward the intended guide strand^11^. The mature guide strand sequence and passenger strand design (including 5′ phosphorylation status and 1 mm position) are shown in figures 1B and 6A.

**Figure 1.**
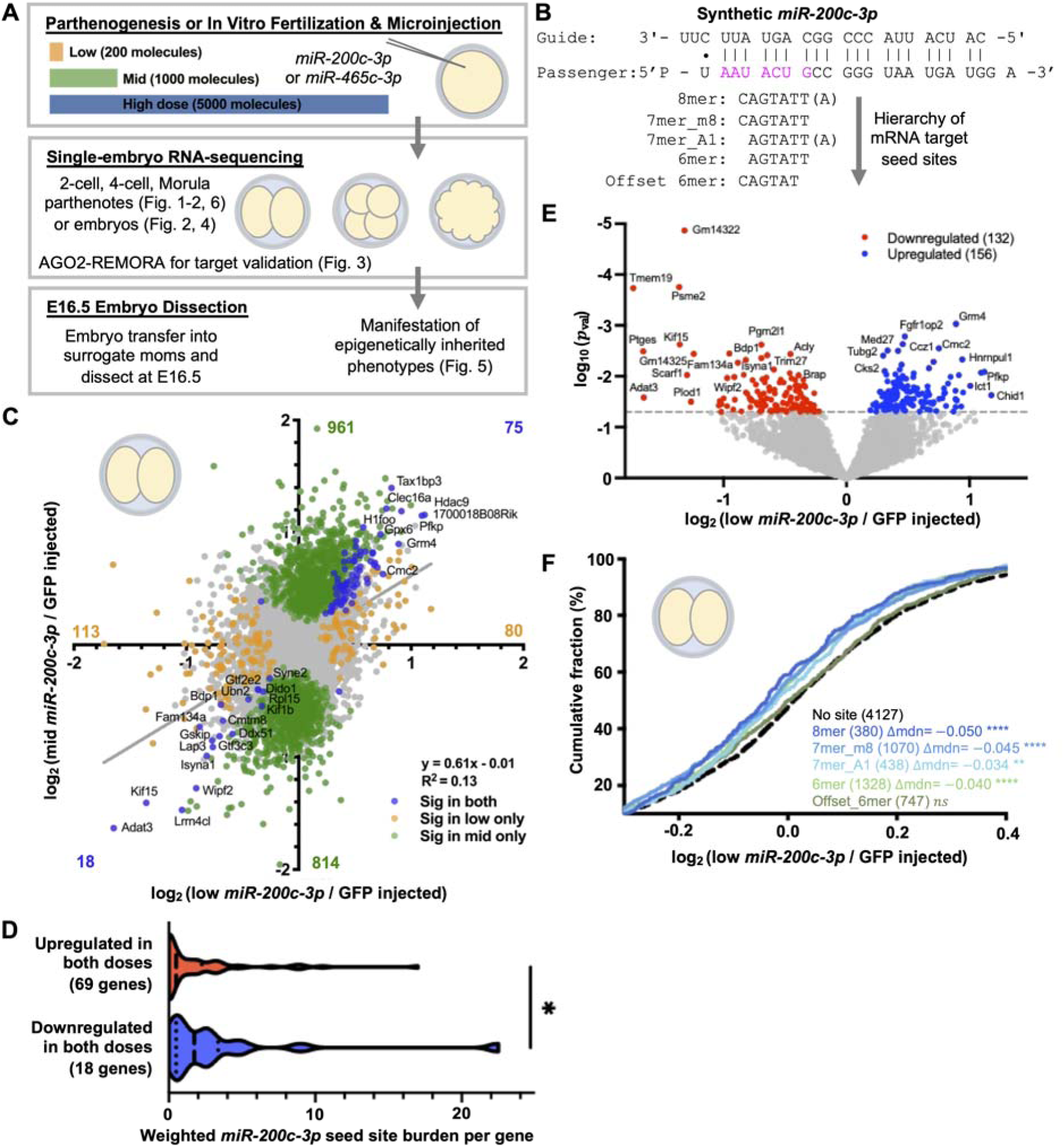
Physiologic *miR-200c-3p* levels induce dose-dependent, seed-sequence directed gene expression regulation in 2-cell embryos. **(A)** Overall experimental schematic of this research. Parthenogenetic or IVF embryos were microinjected with synthetic *miR-200c-3p* or *miR-465c-3p* at low (200 molecules), mid (1000 molecules), or high (5000 molecules) doses. Single-embryo RNA-sequencing was performed at the 2-cell, 4-cell, and morula stages. AGO2- REMORA was used at the 2-cell stage for target identification, and embryos were later assessed for epigenetically inherited phenotypes at E16.5. **(B)** Design of synthetic RNA engineered to preferentially load *miR-200c-3p*. **(C)** Correlation of log_2_ fold change (DESeq2) between low-dose (x-axis) and mid-dose (y-axis) *miR-200c-3p* injection relative to GFP–injected controls in 2-cell parthenotes. Genes with *p* < 0.05 are highlighted as indicated in the legend. **(D)** Distribution of weighted *miR-200c-3p* seed-site burden per gene among genes significantly upregulated or downregulated in both low- and mid-dose 2-cell parthenotes. Statistical significance was assessed using a Mann-Whitney test. **(E)** Volcano plot of differential gene expression in 2-cell parthenotes following low-dose *miR-200c-3p* injection relative to GFP–injected controls, as identified by DESeq2. Genes with *p* < 0.05 are highlighted as indicated in the legend. **(F)** Empirical cumulative distribution function (ECDF) of log_2_ fold changes in low-dose–injected 2-cell parthenotes relative to controls, stratified by mutually exclusive *miR-200c-3p* seed-site class within 3’ UTRs. Distributional shifts relative to genes lacking canonical seed sites were tested using Wilcoxon rank-sum tests. Median differences relative to no-canonical-seed baseline are reported. Asterisks denote significance *(*p* < 0.05), **(*p* < 0.01), ***(p< 0.01), ****(*p* < 0.0001).

### Duplex formation and handling

Lyophilized oligonucleotides were resuspended in nuclease-free water. To generate miRNA duplexes, guide and passenger strands were combined at equimolar concentrations and annealed by incubation at 25°C for 10 minutes. Working and stock solutions were aliquoted and stored at -80°C to minimize freeze-thaw cycles.

### Dose preparation

Duplex miRNAs were generated by serial dilution to the indicated final concentrations for injection and stored in aliquots to ensure replicates received roughly identical doses. miRNA concentrations for microinjection were low = 0.093 ng/µL, mid = 0.46 ng/ µL, high = 2.3 ng/µL. Assuming an injection volume of ∼53 fL, these concentrations correspond to approximately 200, 1000, and 5000 molecules per embryo. GFP-His3.3 mRNA (∼50 ng/µL) encoding green fluorescent protein fused to the histone variant H3.3 was used as an injection control for all injection experiments except AGO2-REMORA. Not all dose conditions were tested at every developmental stage for both miRNAs. Dose selection was guided by prior experimental results and focused on the dose ranges that were most informative at each stage.

### Embryo Microinjection

Cytoplasmic microinjections were performed on parthenotes or embryos as described previously^9^, delivering ∼53 fL of injection solution per embryo. For each experimental replicate, embryos were collected from ∼5 females derived from one or two litters, pooled, and randomly assigned to all injection conditions to minimize litter-specific effects. Within each experimental day, all treatments groups were represented in parallel. Embryos were distributed to ensure approximately matched numbers per condition (typically ∼8 embryos per group and embryonic stage per replicate), preventing confounding by collection date or maternal origin. At least 24 embryos per condition were collected across three independent biological replicates. In one experiment with high embryo yield and injection efficiency, sufficient embryos were obtained across two replicates to meet this threshold.

### AGO2-REMORA Plasmid Construction and In Vitro Transcription

The RNA Adenosine Base Editor (rABE) developed by the Floor laboratory was codon-optimized for *Mus musculus* and synthesized to generate fusion constructs (Twist Bioscience). rABE was fused to either AGO2 or mRUBY3 via a flexible peptide linker (DSGGSSGGSSGSETPGTSESATPESSGGSSGGS) positioned at the C-terminus of rABE. The fusion coding sequences were subcloned into the pRN3P.H3.3-GFP backbone by replacing H3.3-GFP between SpeI and BstII restriction sites. The resulting plasmids, pRN3P.rABE-AGO2 and pRN3P.rABE-mRUBY3, retained the vector’s T3 promoter, Xenopus globin 5′ untranslated region (UTR), 3′ UTR, and poly(A)_30_ tail. A Kozak consensus sequence was introduced into both constructs using the Q5 Site-Directed Mutagenesis Kit (NEB). Plasmids were propagated in *E. coli* NEB 10-beta, purified using the PureLink HiPure Plasmid Midiprep Kit (Thermo Fisher Scientific), and validated by whole-plasmid sequencing (Plasmidsaurus). For in vitro transcription (IVT), plasmids were linearized and transcribed with the mMESSAGE mMACHINE T3 Transcription Kit (Thermo Fisher Scientific) according to the manufacturer’s protocol. IVT products were purified by phenol-chloroform extraction followed by isopropanol precipitation, and RNA integrity was confirmed by agarose gel electrophoresis.

rABE-mRUBY3 mRNA was injected alone to quantify background adenosine deaminase activity. For identification of AGO2-associated transcripts, rABE-AGO2 mRNA was injected either alone or in combination with a synthetic miRNA duplex. Purified mRNAs were adjusted to equivalent molar concentrations (208 nM) prior to injection.

### Single-embryo RNA-sequencing

Following microinjection, individual embryos were collected 28 hours (late 2-cell), 46 hours (4-cell), or 70 hours (compacted morula) post activation or fertilization in 5 μL of TCL buffer (Qiagen, Cat. No. 1070498) with 1% 2-mercaptoethanol. Samples were immediately placed in -80°C until all replicates were collected. mRNA-se libraries were generated using the SMART-Seq protocol^12^. Briefly, lysed embryos were isolated using RNAClean-XP beads and polyadenylated RNA was reverse transcribed using Superscript II. cDNA was amplified and indexed using the Nextera XT kit and sequenced on an Illumina

NextSeq 1000/2000 platform (paired-end 50 bp, ∼5 million reads per sample). Reads were mapped against the *Mus musculus* (mm10) genome using RSEM.

### RNA-sequencing Analysis

To exclude low-complexity libraries, single embryos were required to exhibit detectable expression (≥1 read) in a minimum number of genes. Thresholds were selected separately for each developmental stage and dataset based on inspection of the distribution of detected genes per sample, using the inflection point preceding the major drop-off in library complexity. Stage-specific cutoffs ranged from 6000-8000 detected genes. Samples failing these criteria were excluded from downstream analyses. Raw gene-level counts were analyzed in R (v4.5.1) using DESeq2^13^ (v1.48.2). Genes were retained if they achieved at least 2.5 counts per million (CPM) in at least 80% of samples within at least one treatment group. Genes failing these criteria were removed from downstream analysis. Differential expression was modeled using a negative binomial generalized linear model with batch (embryo collection date) as a variable and Wald tests were performed to assess differential expression between each treatment level and designated control group. Normalized counts and full differential expression statistics were exported for downstream analysis. Given the limited statistical power of single-embryo sequencing due to the nature of the experimental design, nominal p-values were used for candidate identification, while enrichment and pathway-level analyses incorporated multiple-testing correction.

### Identification of miRNA Seed Sites in 3’ UTRs

Mouse 3’ UTR sequences were derived from the GENCODE mm10 annotation (gencode.vM10.gtf). All annotated transcripts containing 3’ UTRs were retained. Canonical seed sites were defined using nucleotides 2-8 of the mature miRNA sequence. Reverse-complement motifs corresponding to 8mer, 7mer_m8, 7mer_A1, 6mer, offset 6mer (positions 3-8) were scanned across 3’ UTR sequences using Biostrings. As a representative noncanonical seed match, 7mer_1mm sites were defined as 7 nucleotide matches containing exactly one mismatch relative to the canonical 7mer_m8 seed (positives 2-8). Genes were annotated for presence or absence of each site class. For analyses requiring assignment of a single site class per gene, a hierarchical classification was applied prioritizing stronger canonical sites (8mer > 7mer_m8 > 7mer_A1 > 6mer > offset 6mer), with genes lacking canonical sites classified separately.

### Enrichment and Distribution Analyses

Enrichment of seed sites among gene sets was assessed using two-sided Fisher’s exact tests comparing genes with at least one site of a given class to all other genes with annotated 3’ UTRs. Odds ratios and p-values were calculated.

To assess the relationship between site class and gene expression changes, Empirical cumulative distribution frequencies (ECDFs) of log_2_ fold changes were generated. Analyses were performed using mutually exclusive (hierarchical strongest-site assignment) with genes lacking canonical sites serving as the baseline distribution unless otherwise indicated.

### Computational Analysis for AGO2-REMORA

Raw sequencing reads were assessed for quality, including adapter content, using FastQC. Reads were aligned to the Mus musculus GENCODE VM25 annotation (mm10-compatible) using STAR (v2.7.10a)^14^, allowing up to 10 multiple mapping loci per read (-- outFilterMultimapNmax 10), with a maximum of 10 mismatches per read (-- outFilterMismatchNmax 10), and no more that 30% mismatches relative to read length (-- outFilterMismatchNoverLmax 0.3), and including the MD tag in output alignments.

Mapped reads were processed to mark PCR duplicates using Picard Tools MarkDuplicates (v2.26.10, Broad Institute). Alignments were further filtered to retain only properly paired reads with mapping quality >20, following prior SAILOR pipeline guidelines.^15^ Filtered BAM files were used as input to the MARINE pipeline (Yeo Lab), restricting analysis to standard chromosomes and specifying the –strandedness parameter to account for unstranded libraries.

Identified editing sites were annotated to gene regions using the corresponding GENCODE VM25 annotation BED file. Sites overlapping known single nucleotide polymorphisms were removed based on dbSNP (mm10 build). SNP filtering was performed in R (v4.4.0) using the Bioconductor packages GenomicRanges (c1.56.2) and rtracklayer (v1.64.0).

### RNA editing quantification and normalization

A-to-G editing counts were aggregated by summing site-levels counts within each gene and genomic region (coding sequence, 3’ UTR, or intron) for each sample. Region definitions and gene/region lengths were derived from the GENCODE mouse GTF (gencode.vM10). Transcript-level intervals were collapsed to gene-level unions by gene symbol. Intronic intervals were defined per gene as the gene body minus the union of annotated exons. For each sample, the number of RNA-sequencing (RNA-seq) alignments overlapping coding sequence (CDS), inferred 3’ UTRs, or introns were quantified from genome-aligned BAM files using featureCounts in SAF mode, producing region-specific mapped-read denominators. Editing activity was normalized to region length and region-specific library depth by computing editing events per kilobase per million mapped reads (EPKM); EPKM = edit_counts x10^9^/(mapped_reads_region_ xregion_bp). The resulting gene-by-sample EPKM matrices for CDS, 3’ UTR, and introns were used for downstream comparisons across experimental conditions.

### Statistical identification of miRNA-directed RNA editing targets

To account for small differences in global editing activity across conditions given the nature of embryo microinjections, EPKM values were further normalized within each replicate by scaling to a total of one million, generating edits per million (EPM) values that represent the fraction of total editing attributable to each transcript. Per-gene RNA editing levels were quantified for each single embryo as editing events per million mapped reads (EPM). To stabilize variance and accommodate zero-inflated distributions, EPM values were transformed at the embryo level as log₂(EPM + 1). For each gene, the mean log₂(EPM + 1) was calculated separately across rABE_Ago embryos and rABE_Ago + miRNA embryos. The change in editing was defined as the difference in these means. Editing abundance was defined as the average of the two group means. These values were visualized in an MA-style plot to separate editing abundance from *miR-200c-3p* and *miR-465c-3p* induced editing shifts.

Statistical significance was assessed per gene using an unpaired one-sided Wilcoxon rank-sum test comparing log₂(EPM+1) values between conditions. Genes were required to have data from at least three embryos per group to be tested. Given the limited power and inherent variability of single-embryo RNA-sequencing, nominal *p*-values were interpreted alongside effect size and reproducibility across embryos.

### Gene Functional Classification

Genes were functionally annotated using the Ensembl BioMart interface. Gene identifiers were queried against the *Mus musculus* dataset using the GRCm39 genome assembly to retrieve associated Gene Ontology (GO) annotations. Genes were classified into the following categories: Transcription factors (TFs): GO:003700, GO:0000981, GO:0001228, GO:0001227, GO:0000977, GO:0000978, GO:0000987, GO:0140297; RNA-binding protein (RBP): GO:0003723, GO:0003729, GO:0044822; Chromatin-associated: GO:0000785, GO:0003682, GO:0006325, GO:0006338. Genes annotated in multiple categories were assigned hierarchically, prioritizing TFs, chromatin-associated, then RBPs.

### Ingenuity Pathway Analysis and Canonical Pathway Visualization

Ingenuity Pathway Analysis (IPA; QIAGEN) was performed on differentially expressed genes from 4-cell and morula-stage sperm-fertilized embryos injected with *miR-200c-3p*. Canonical pathways were filtered based on minimum gene overlap (≥5 genes for 4-cell and ≥10 genes for morula) to account for stage-specific differences in transcriptional complexity. To reduce redundancy among highly overlapping pathways, gene sets were clustered using Jaccard similarity (threshold = 0.3), and connected components were identified. For each cluster, a single representative pathway was selected based on lowest p-value, followed by absolute z-score and number of overlapping genes.

### E16.5 Embryo Generation and Collection

ICR Females (Envigo) were paired overnight with vasectomized males, and the following morning females were assessed for copulation plugs. Plugged females were separated and used as recipients for surgical embryo transfer. IVF embryos injected with low-dose *miR-200c-3p* with GFP or GFP-only controls were transferred to pseudopregnant females. At embryonic day (E) 16.5, pregnant females were euthanized and dissected to obtain fetuses.

### Digital image acquisition and processing

During dissections, we collected digital photographs of the front, left, and right profiles of each fetus within each litter. We then processed the images for morphometric analyses using methods described previously^7,16^. Briefly, we imported digital images of the facial profiles into the publicly available image analysis software tpsDig2w64^17^ (version 2.32). We set the reference scale bar in the picture to 1 mm and then demarcated the eighteen facial landmarks described by Anthony et al. (2010)^18^. To ensure consistency, a single individual (S.L.H.) demarcated the landmarks in each photograph, consistently identifying the exact location and order for each image. We then created a linear outline around the head and digitized the landmarks and outlines as a T.P.S. file. We then used the publicly available program tpsDig2w64^17^ to add additional landmarks, including the midpoints between features and other aspects of the outline, producing a total of 47 landmarks. In curating this dataset, we named each file with the litter ID, sex, and uterine position of each fetus and then used the publicly available program tpsUtil32^19^ (version 1.83) to generate a T.P.S. database.

### Geometric Morphometrics and Statistical Analyses of Facial Images

We imported the generated T.P.S. files for each fetus into the MorphoJ software^20^(version build 1.07a, Java version 1.8.0_291 (Oracle Corporation)) and conducted geometric morphometric analysis using methods described previously^7,16^. Briefly, we added classifiers describing each treatment group and then separately normalized the datasets for scale, rotation, and translation using the Procrustes fit feature^20^. We then generated a covariance matrix, which we used to conduct Principal Component Analysis (PCA). We then used Canonical Variate (CV) analysis to identify differences in facial features between treatments and exported the raw CV scores into the publicly available Paleontological Statistics Software Package for Education and Data Analysis (PAST) analysis software^21^ (version 4.03; [https://softfamous.com/postdownload-file/past/18233/13091/.]). We conducted multivariate analyses of the raw CV scores using statistical methods described previously^22–25^. These included the parametric Multivariate analysis of variance (MANOVA) and Nonparametric Analysis of similarities (ANOSIM), and Permutational multivariate analysis of variance (PERMANOVA) tests, followed by Bonferroni correction. We generated the CV wireframe and scatter plots using the graphing features of MorphoJ^20^.

### Figure generation

Graphs and statistical analysis for Figures 1-3, 4A, 4C, Figure 6 and Supplemental Figures were created using Prism (v10.7.0) and final edits were completed in PowerPoint. Figures 4B and 4D were generated in R (v4.5.1).

## RESULTS

Parthenogenetic embryos (parthenotes) are chemically activated eggs that undergo preimplantation embryonic development in the absence sperm^26^. This system enables direct interrogation of individual sperm RNAs in isolation from the entire sperm RNA payload. To determine whether modest changes in sperm miRNA abundance are sufficient to overcome dilution within the much larger egg, we first established a physiologically calibrated perturbation range. A single mouse sperm is estimated to carry hundreds of thousands of miRNA molecules^27^ with a subset (∼40) accounting for most of the total miRNA pool (Supplemental Table 1). Based on this distribution, we conservatively estimated hundreds of *miR-200c-3p* molecules are normally delivered to the zygote by sperm (Supplemental Figure 1A). Prior studies demonstrating causal roles for sperm miRNAs in the transmission of non-genetically inherited phenotypes have typically used a concentration of ≥1 ng/µL of synthetic miRNA, corresponding to about 2000 molecules of miRNA per zygotic injection^3,4,6^. Therefore, to better define the dynamic range over which sperm miRNAs influence embryonic gene regulation, we also tested beyond the endogenous physiologic range to include supraphysiologic doses (5000 molecules). This approach allowed us to assess dose-responsiveness and potential saturation effects within the early embryo.

### *miR-200c-3p* drives dose-dependent repression at the 2-cell stage

To define the immediate gene expression consequences of altered sperm miRNA abundance, we microinjected zygotes with defined quantities of *miR-200c-3p* spanning the estimated endogenous delivery range (low, 200 molecules), a scaled upper physiologic level commonly used in function studies (mid, 2000 molecules), and a higher supraphysiologic dose (high, 5000 molecules), and collected embryos at successive developmental stages for transcriptomic analysis (Figure 1A). We first focused on the late 2-cell stage (28-hours after parthenogenetic activation), shortly following zygotic genome activation, to capture the earliest regulatory responses to miRNA perturbation. To isolate strand-specific activity, we used synthetic duplexes that preferentially load the *miR-200c-3p* guide strand^11^(Figure 1B). Single-embryo RNA sequencing at the 2-cell stage revealed that even the lowest, physiologic dose elicited gene expression changes relative to GFP–controls, with consistent and enhanced effects at the mid-dose (Figure 1C). Further, log_2_ fold changes were positively correlated between low- and mid-doses with R^2^ = 0.13 (Figure 1C). 80% of genes significantly upregulated by the mid-dose exhibited a positive log_2_ fold change in the low-dose, and 65% of genes downregulated by the mid-dose exhibited a negative log_2_ fold change in the low-dose. These results indicate a structured regulatory program that scales quantitatively with miRNA input rather than reflecting stochastic or nonspecific RNA effects.

Importantly, genes consistently altered across doses showed selective enrichment for *miR-200c-3p* targeting among downregulated transcripts. The 18 transcripts significantly repressed by both doses carried a significantly higher burden of *miR-200c-3p* seed site matches in their 3’ UTRs than upregulated genes, consistent with direct, sequence-dependent repression of targets (Figure 1D). Extending these observations to the transcriptome more broadly, low-dose *miR-200c-3p* injection produced transcriptional changes relative to GFP–injected controls, with hundreds of genes significantly up- and down-regulated (Figure 1E). Stratifying genes by containing canonical seed-sites revealed systematic negative shifts in log₂ fold-change distributions for transcripts containing *miR-200c-3p* sites in their 3′ UTRs, with stronger shifts observed for higher-affinity site types (Figure 1F). Consistent with prior studies demonstrating that noncanonical interactions also contribute to miRNA target recognition,^28,29^ we observed that genes harboring a single mismatch within a 7mer seed also exhibit downregulation compared genes lacking either canonical or mismatch sites (Supplemental Figure 1G).

Parallel experiments using *miR-465c-3p* produced a distinct gene expression response at both low- and high-doses, demonstrating that different sperm miRNAs have unique regulatory effects throughout early embryonic development (Supplemental Figure 1B-F). Together, these results demonstrate that physiologically relevant perturbations to a single sperm miRNA, *miR-200c-3p,* are sufficient to drive coordinated, sequence-directed repression programs at the onset of zygotic genome activation consistent with direct miRNA-mediated regulatory activity.

### Tunable miRNA-dependent gene regulation unfolds across preimplantation development

We next asked whether the initial regulatory effects of *miR-200c-3p* are transient, persistent, or modulated as development proceeds. To examine how miRNA-induced regulatory programs unfold across subsequent developmental stages, we analyzed gene expression at the 4-cell and morula stages following zygotic injection of our calibrated titration series. At both stages, log₂ fold changes were strongly positively correlated across low-, mid-, and high-doses, demonstrating gene expression responses scale quantitatively with increasing miRNA input rather than diverging at higher doses (Figure 2A-B, Supplemental Figure 2A-D).

**Figure 2.**
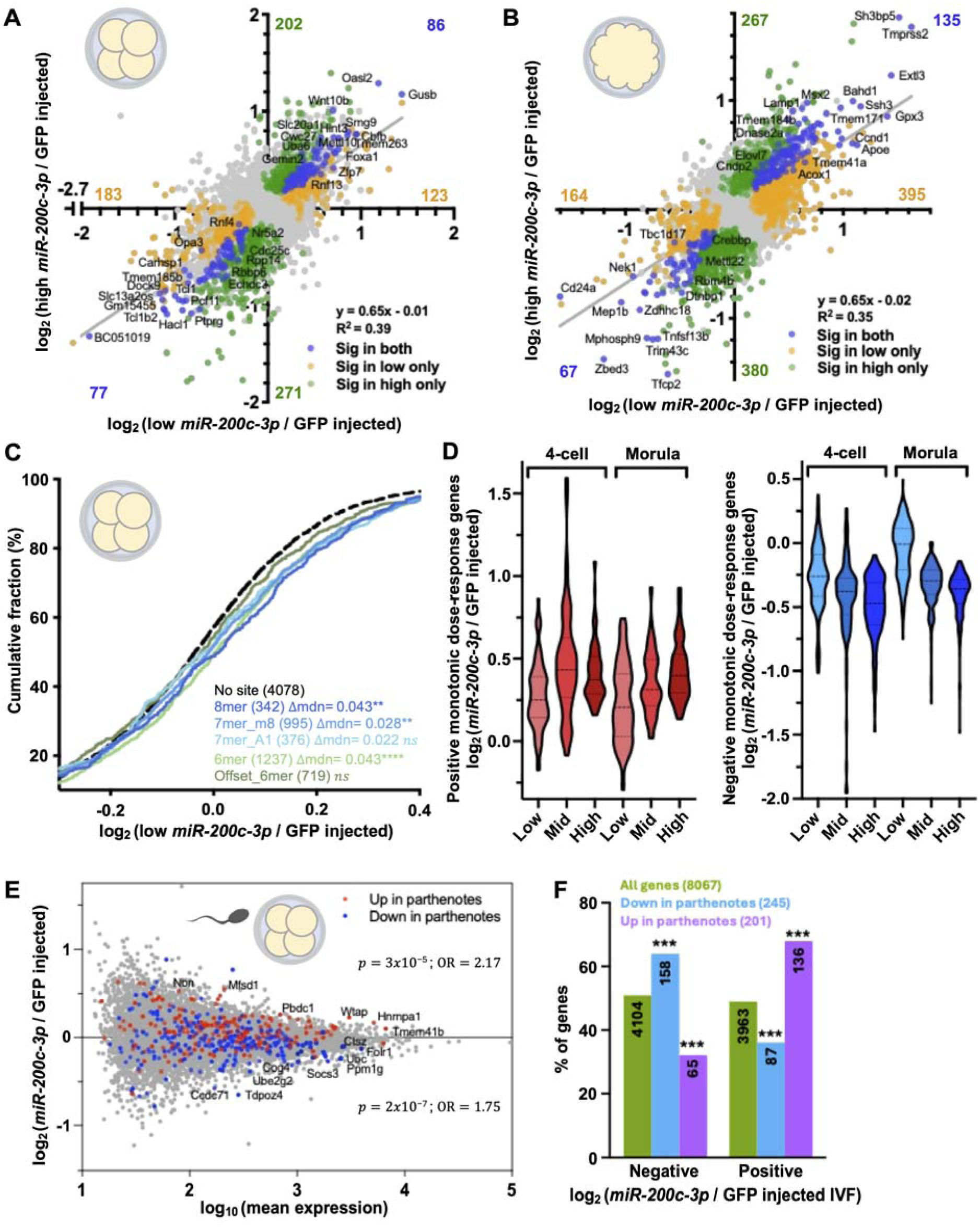
Tunable, dose-dependent transcriptional responses initiated by *miR-200c-3p* propagate across preimplantation development. **(A-B)** Correlation of log_2_ fold change (DESEq2) between low (x-axis) and high-dose (y-axis) *miR-200c-3p* injection relative to GFP– injected controls in 4-cell (A) and morula (B) parthenotes. Genes with *p* < 0.05 are highlighted as indicated in the legend. **(C)** ECDF of log_2_ fold changes in 4-cell parthenotes for genes stratified by mutually exclusive *miR-200c-3p* seed site class in 3’ UTRs following low-dose injection. Distributional shifts relative to genes lacking canonical seed sites were tested using Wilcoxon rank-sum tests. Median differences relative to no-canonical-seed baseline are reported. **(D)** Distribution of log_2_ fold changes (DESeq2) for genes exhibiting positive (left) or negative (right) monotonic dose-response behavior across low-, mid-, and high-dose *miR-200c-3p* injection at the 4-cell and morula stages. Dose-dependent genes were identified using a likelihood ratio test (LRT; nominal *p* < 0.05), and the direction of dose response was classified using a numeric-dose Wald test. n = 88 activated and 124 repressed genes in 4-cells; n = 88 activated and 219 repressed genes in morula. **(E)** MA plot of gene expression changes at the 4-cell stage in *miR-200c-3p*-injected sperm-fertilized IVF embryos relative to GFP-injected controls. Genes significantly dysregulated in 4-cell parthenogenetic embryos following low-dose *miR-200c-3p* injection are highlighted. Fisher’s exact test statistics, odds ratios (OR) and two-sided p - values for up- and down-regulated genes in parthenotes are shown. **(F)** Fisher’s test enrichment of genes up- or downregulated in parthenotes among genes exhibiting positive or negative log_2_ fold changes in IVF embryos following *miR-200c-3p* injection. Asterisks denote significance *(*p* < 0.05), **(*p* < 0.01), ***(*p* < 0.001), ****(*p* < 0.0001).

Strikingly, the directionality of target gene behavior shifted across stages. At the 4-cell stage, transcripts harboring canonical *miR-200c-3p* seed sites showed increased expression relative to controls following low-dose injection in contrast to their repression at the 2-cell stage (Figure 2C). Notably, 80% of the 3,201 genes containing canonical seed sites at the 2-cell stage are expressed and globally upregulated at the 4-cell stage. This stage-specific reversal is consistent with a rebound or compensatory increase in target transcripts that were initially repressed. Correspondingly, significantly up- and down-regulated genes were enriched for seed sites (Supplemental Figure 2E). This pattern was more pronounced at the mid-dose (Supplemental Figure 2F), indicating that a modestly increased miRNA input is associated with a stronger stage-specific shift in target gene expression. Unexpectedly, a partial attenuation of seed-dependent repression was already detectable at the 2-cell stage under mid-dose conditions (Supplemental Figure 1H), suggesting that incremental changes in miRNA input are sufficient to fine-tune downstream transcriptional responses temporally. By the morula stage, this seed-dependent bias was no longer evident (Supplemental Figure 2G), consistent with a model in which early direct targeting initiates a transcriptional cascade that becomes uncoupled from direct seed-site enrichment at later stage.

To formally quantify dose responsiveness at each stage, we modeled gene expression as a function of miRNA dose using a likelihood ratio test (LRT). At the 4-cell stage, 88 genes showed progressive activation and 124 showed progression repression with increasing miRNA dose. By the morula stage, 88 and 219 genes, exhibited monotonic dose-dependent activation and repression, respectively (Figure 2D, Supplemental Figure 2H-I). These results demonstrate that *miR-200c-3p* induces gene expression changes that persist across preimplantation development and increase in magnitude with higher miRNA input. While direct, seed-dependent repression predominates at the 2-cell stage, later stages exhibit broader transcriptional changes that scale with miRNA dose but are no longer restricted to canonical seed-site targets.

We next asked whether the gene expression changes identified in parthenogenetic embryos following *miR-200c-3p* are recapitulated in sperm-fertilized embryos. In an equivalent experiment, we injected a low- dose of *miR-200c-3p* into in vitro fertilized (IVF) embryos to model a modest increase in *miR-200c-3p* abundance within the context of fertilization. Genes upregulated in parthenotes were significantly enriched among transcripts exhibiting positive log₂ fold changes in *miR-200c-3p*–injected IVF embryos, whereas genes downregulated in parthenotes were preferentially enriched among transcripts exhibiting negative fold changes (Figure 2E). Parthenote-upregulated genes were 2.17-fold more likely to exhibit positive values compared to background genes, while downregulated genes were 1.76-fold more likely to exhibit negative values. Furthermore, genes downregulated in parthenotes were depleted for positive fold changes in sperm-fertilized embryos, whereas genes upregulated in parthenotes were depleted for negative fold changes in embryos (Figure 2F). An even stronger concordance was observed when using the mid-dose–responsive genes to make the same comparison (Supplemental Figure 2J). These findings indicate that *miR-200c-3p*–dependent transcriptional responses identified in parthenotes are not artifacts of the parthenogenetic system but instead reflect regulatory programs relevant to natural embryonic development.

### Direct targets of *miR-200c-3p* identified in preimplantation embryos using AGO2-REMORA

Direct identification of miRNA targets in preimplantation embryos has remained technically challenging due to the limited material available at early embryonic stages precluding conventional biochemical approaches such as Argonaute immunoprecipitation. To distinguish direct *miR-200c-3p* targeting from secondary transcriptional responses, we adapted RNA-encoded molecular recording in adenosines (REMORA)^30^ by fusing an optimized RNA adenosine base editor, rABE, to Argonaute2 (AGO2-REMORA). This strategy enables site-resolved A-to-I editing at AGO2-bound transcripts, thereby creating a molecular footprint of miRNA-dependent interactions in vivo while controlling for baseline editing and AGO2 activity.

We performed single-embryo RNA-seq at the late 2-cell stage following injection of rABE alone, rABE_AGO2, or rABE_AGO2 plus a low-dose of synthetic miRNA (Figure 3A). Editing activity increased in 3’ UTRs and coding regions upon AGO2 tethering, while intronic editing remained comparable to baseline levels, consistent with selective AGO2 engagement of mature mRNAs rather than global editing artifacts (Figure 3B, Supplemental Figure 3A). Editing was not further altered by the addition of *miR-200c-3p* or *miR-465c-3p* indicating that changes in editing reflect redistribution of AGO2 occupancy rather than global increases in editing activity.

**Figure 3.**
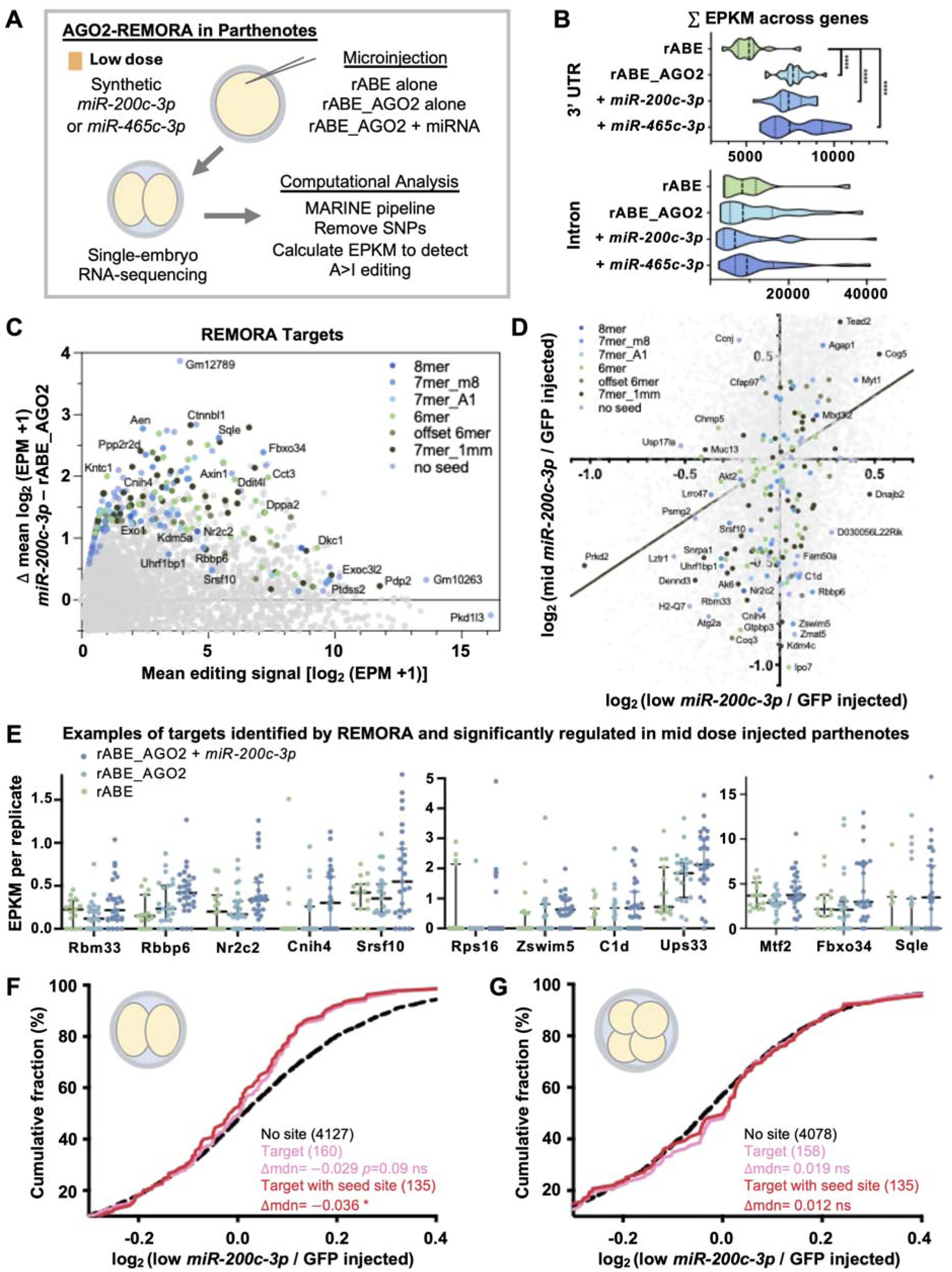
AGO2-REMORA identifies direct *miR-200c-3p* target mRNAs in early embryos. **(A)** Experimental schematic for AGO2-REMORA target identification. Parthenogenetic embryos were microinjected with mRNA constructs encoding rABE alone, rABE_AGO2, or rABE_AGO2 plus synthetic *miR-200c-3p* or *miR-465c-3p*. Single-embryo RNA sequencing was performed to quantify gene-level RNA editing events as a proxy for AGO2 engagement. **(B)** Distribution of summed editing events per kilobase mapped reads (EPKM) per replicate in 2-cell parthenotes across experimental conditions in 3’ UTRs (upper) and introns (lower). **(C)** MA plot showing gene level editing abundance in low-dose 2-cell parthenotes. The x-axis shows the average log_2_(EPM+1) across rABE_AGO2 and rABE_AGO2+*miR-200c-3p* parthenotes, and the y-axis represents the difference in average log2(EPM+1) between conditions (rABE_AGO2+*miR-200c-3p* minus rABE_AGO2). Genes exhibiting significantly increased editing upon *miR-200c-3p* addition were identified using a one-sided Wilcoxon rank-sum test on embryo-level log_2_(EPM+1) values and are highlighted as indicated in the legend. Highlighted genes have nominal *p* < 0.05**. (D)** Genes exhibiting significant *miR-200c-3p*–dependent increases in editing (*miR-200c-3p* targets) were overlaid onto the 2-cell parthenote RNA-seq dose-response correlation plot from Figure 1C. **(E)** Gene-level editing signal (EPKM) in 2-cell parthenotes for target mRNAs identified by AGO2-REMORA containing canonical 8mer or 7mer seed sites and downregulated by mid-dose *miR-200c-3p* injection at the 2-cell stage. Median with 95% confidence intervals are indicated. Y-axis scales differ between panels. **(F-G)** ECDF of log_2_ fold-changes (DESeq2) in 4-cell (F) and morula (G) following low-dose *miR-200c-3p* injection in parthenotes compared to control for target mRNAs identified by AGO2-REMORA. Distributional shifts relative to genes lacking canonical seed sites were tested using Wilcoxon rank-sum tests. Median differences relative to no-canonical-seed baseline are reported. Asterisks denote significance *(*p* < 0.05), **(*p* < 0.01), ***(*p* < 0.001), ****(*p* < 0.0001).

To identify AGO2-REMORA-defined direct targets, we quantified gene level editing differences between rABE_AGO2 and rABE_AGO2 + *miR-200c-3p* parthenotes. We identified 207 genes exhibiting significant *miR-200c-3p*–dependent increases in editing relative to controls (Figure 3C). Among these, 134 genes contained either canonical or noncanonical *miR-200c-3p* seed sites. The complete distribution of editing abundance and *miR-200c-3p*–dependent shifts across all transcripts is shown in Supplemental Figure 3B.

Analysis of AGO2-REMORA editing-defined targets with 2-cell transcriptomic data (Figure 1C) revealed substantial concordance between AGO2 engagement and dose-dependent transcriptional repression. Of the 207 editing-enriched genes, 63 exhibited negative log_2_ fold changes at both low- and mid-doses, while an additional 47 were repressed at the mid-dose. In contrast, only 35 genes were positively regulated at both doses (Figure 3D). Consistent with these trends, 7 of 1,037 genes significantly upregulated at mid-dose were identified as targets, whereas 37 of 833 downregulated were editing targets, indicating preferential overlap between increased AGO2 occupancy and transcriptional repression at the 2-cell stage. AGO2-REMORA defined targets containing canonical 7mer or 8mer seed sites and significantly downregulated in the mid-dose *miR-200c-3p* RNA-seq dataset exhibited increased AGO2-associated editing following *miR-200c-3p* addition (Figure 3E).

Furthermore, cumulative distribution analysis of low-dose 2-cell embryo RNA-seq log_2_ fold change demonstrated a negative expression shift among AGO2-REMORA defined targets, which became more pronounced and statistically significant when restricting to targets harboring seed sites (Δmedian = -0.036, p < 0.05) (Figure 3F). Consistent with a dose-dependent strengthening of direct targeting, AGO2-REMORA defined targets exhibited significantly stronger negative log_2_ fold changes at the 2-cell stage under mid-dose conditions relative to low-dose (Supplemental Figure 3D). In contrast, this association was not observed at the 4-cell stage, paralleling the developmental attenuation of seed-dependent repression described above (Supplemental Figure 3E). Notably, many AGO2-REMORA defined direct *miR-200c-3p* targets encode regulatory proteins (annotated by Ensembl) including 14 transcription factors, 14 RNA-binding proteins, and 13 chromatin-associated factors, positioning early miRNA-mediated repression to influence broader developmental gene networks (Supplemental Figure 3E). Together, these results provide direct molecular evidence that *miR-200c-3p* recruits AGO2 to specific endogenous embryonic transcripts in a seed-sequence-dependent manner and establishes a mechanistic link between miRNA induction, AGO2 occupancy, and direct transcript repression during early embryogenesis.

### Stage-specific transcriptional programs emerge following early low-dose *miR-200c-3p* targeting

To assess how *miR-200c-3p*–perturbation impacts broader developmental programs, we performed RNA-seq analyses across later preimplantation embryonic stages in *miR-200c-3p*–injected sperm-fertilized embryos. RNA-seq analysis revealed stage-specific transcriptional responses to *miR-200c-3p* injection. At the 4-cell stage, 161 genes were upregulated and 184 were downregulated relative to controls (Figure 4A). Redundancy-collapsed pathway analysis identified enrichment of DNA repair programs, histone modification signaling, neddylation, and RHO GTPase–associated pathways among others (Figure 4B). By the morula stage, differential expression expanded to 246 upregulated and 375 downregulated genes (Figure 4C). Enriched pathways were different from those observed at the 4-cell stage and included cytoskeletal polarity and trafficking networks, with strong enrichment of Ephrin B signaling, RHO GTPase cycle, protein sorting and Golgi trafficking pathways, as well as chromatin modifying enzymes and metabolic regulatory pathways including sirtuin signaling (Figure 4D). Although some pathway themes were shared between stages, the specific genes driving those enrichments were largely distinct, indicating that transcriptional responses evolve as development proceeds. Together, these results indicate that early *miR-200c-3p* perturbation alters not only individual target transcripts but also produces sustained, stage-specific embryonic changes in gene expression across regulatory networks, providing a mechanistic bridge between low-dose miRNA exposure and long-term developmental phenotypes. Our findings support a ’cascade model’, where sperm miRNAs directly target mRNAs early, initiating downstream gene expression changes that propagate across developmental stages (Figure 4E).

**Figure 4.**
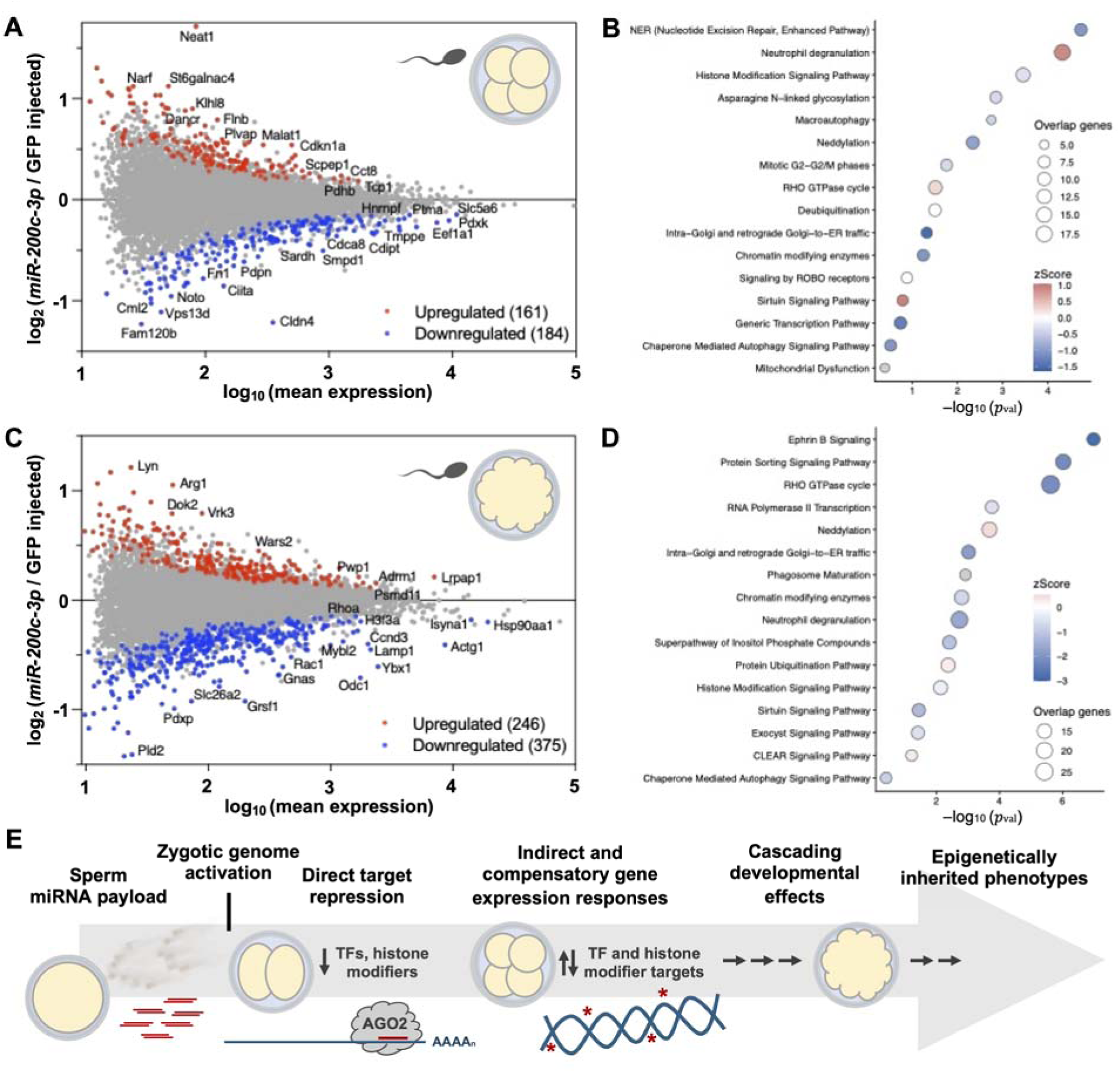
*miR-200c-3p* drives stage-specific gene expression changes in sperm-fertilized embryos. Gene expression analysis (DESeq2) of single embryo RNA-seq data from 4-cell and morula stage embryos injected as zygotes with low-dose *miR-200c-3p*. **(A, C)** MA plots of gene expression changes at the 4-cell **(A)** and morula **(C)** stage embryos following *miR-200c-3p* zygotic low-dose injection. Genes with nominal are highlighted as indicated in the legend. **(B, D)** Ingenuity Pathway Analysis (IPA) of differentially expressed genes at the 4-cell **(B)** and morula **(D)** stages. Canonical pathways are plotted by log_10_ () on the x-axis. Circle size reflects the number of overlapping genes and color indicates IPA z-score. Redundant pathways were collapsed by Jaccard similarity clustering, and a representative pathway per cluster is shown. **(E)** Cascade model illustrating sperm miRNAs directly repressing target transcripts after ZGA followed by secondary and downstream regulation in later stages, ultimately contributing to developmental phenotypes.

### Physiologic *miR-200c-3p* perturbation recapitulates paternal alcohol-associated craniofacial phenotypes

Given prior evidence linking paternal alcohol exposure to elevated sperm *miR-200c-3p* levels and fetal alcohol-associated craniofacial dysmorphology, we next employed a widely used technique for comparing diverse aspects of craniofacial shape, including the characterization of fetal alcohol syndrome-associated facial dysmorphology in both preclinical and clinical studies^31^. We performed geometric analysis on embryonic day 16.5 (E16.5) embryos derived from low-dose *miR-200c-3p*-injected zygotes transferred to surrogate mothers. To quantify craniofacial morphology, we performed geometric morphometric analysis of facial landmarks in E16.5 embryos. Statistical modeling revealed significant variation in facial shape attributable to *miR-200c-3p* exposure. These differences were characterized by altered facial proportions, including changes in centroid size, shifts in eye and ear positioning, and differences in jaw growth in *miR-200c-3p*-injected animals (Supplemental Data 1-3, Figure 5A). Canonical variate analysis revealed clear separation between *miR-200c-3p*-injected and control embryos, indicating distinct craniofacial morphologies (Figure 5B-D). Multivariate statistical testing confirmed significant differences between treatment groups. When stratified by sex, distinct patterns emerged. In males, *miR-200c-3p* injection altered facial shape in the front and left profiles without affecting overall size (Supplemental Data 4-6). In females, facial shape was altered in the left profile and was accompanied by changes in facial size in the front and right profiles (Supplemental Data 7-9). Together, these results demonstrate that modest, physiologically relevant delivery of a single sperm miRNA at the zygote stage is sufficient to induce reproducible, sex-specific craniofacial phenotypes detectable in late-gestation embryos.

**Figure 5.**
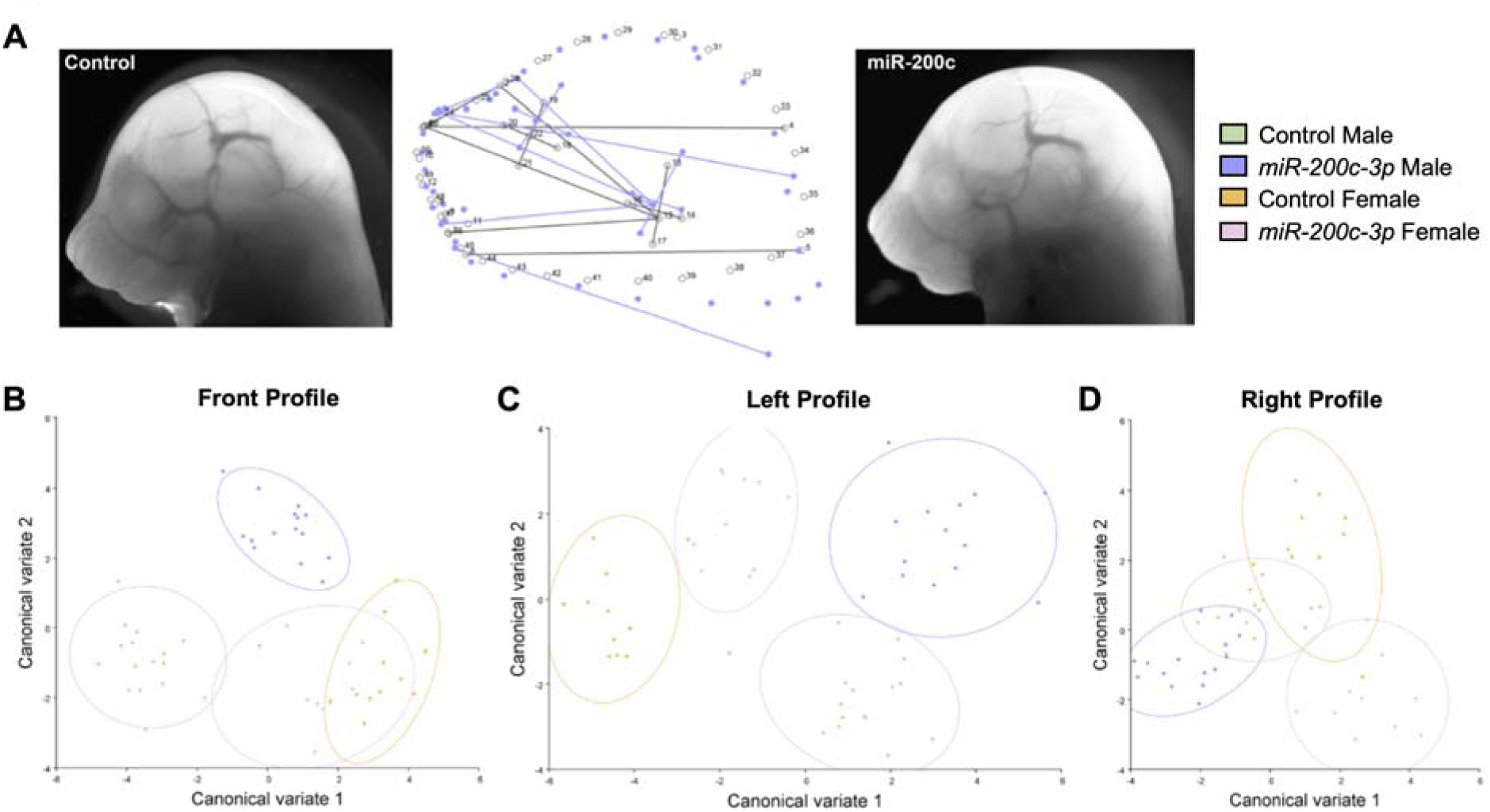
Delivery of 200 molecules of *miR-200c-3p* injection is sufficient to induce sex-specific craniofacial alterations. Zygotes were microinjected with 200 molecules of synthetic *miR-200c-3p* or control and transferred to surrogate females. Embryos were harvested at embryonic day 16.5 (E16.5), and standardized frontal and lateral images were collected for geometric morphometric analysis. Landmark coordinates were aligned using generalized Procrustes analysis to remove differences in scale, orientation, and position prior to statistical comparison of facial shape. **(A)** Representative images of control (left) and *miR-200c-3p*–injected (right) male embryos, flanking a wireframe representation of canonical variate 1 (CV1) illustrating landmark displacement associated with treatment. **(B-D)** Canonical variate plots showing treatment-dependent separation of facial morphology in the front **(B)**, left **(C)**, and right **(D)** profiles. Multivariate analyses confirmed significant differences between treatment groups (see Supplemental Data 1-9 and Methods).

### *miR-465c-3p* elicits a distinct, stage-dependent gene expression program in early embryos

To determine whether additional sperm-enriched miRNAs engage similar or distinct regulatory programs during early development, we performed parallel analyses using synthetic duplexes to preferentially load *miR-465c-3p* (Figure 6A). Parthenogenetic embryos were subjected to a similar dose series (low and high only) and the same analytical framework applied to *miR-200c-3p*. Low-dose *miR-465c-3p* injection into parthenogenetic embryos induced significant transcriptional changes relative to GFP–injected controls, with hundreds of genes differentially expressed at the 2-cell stage (142 downregulated, 73 upregulated; Figure 6B). Similarly to *miR-200c-3p*, transcriptional responses were correlated between low- and high-doses of *miR-465c-3p* at both 4-cell (R^2^ = 0.20) and morula (R^2^ = 0.43) stages, again indicating a coherent regulatory program that scales quantitatively with miRNA input (Figure 6C-D).

**Figure 6.**
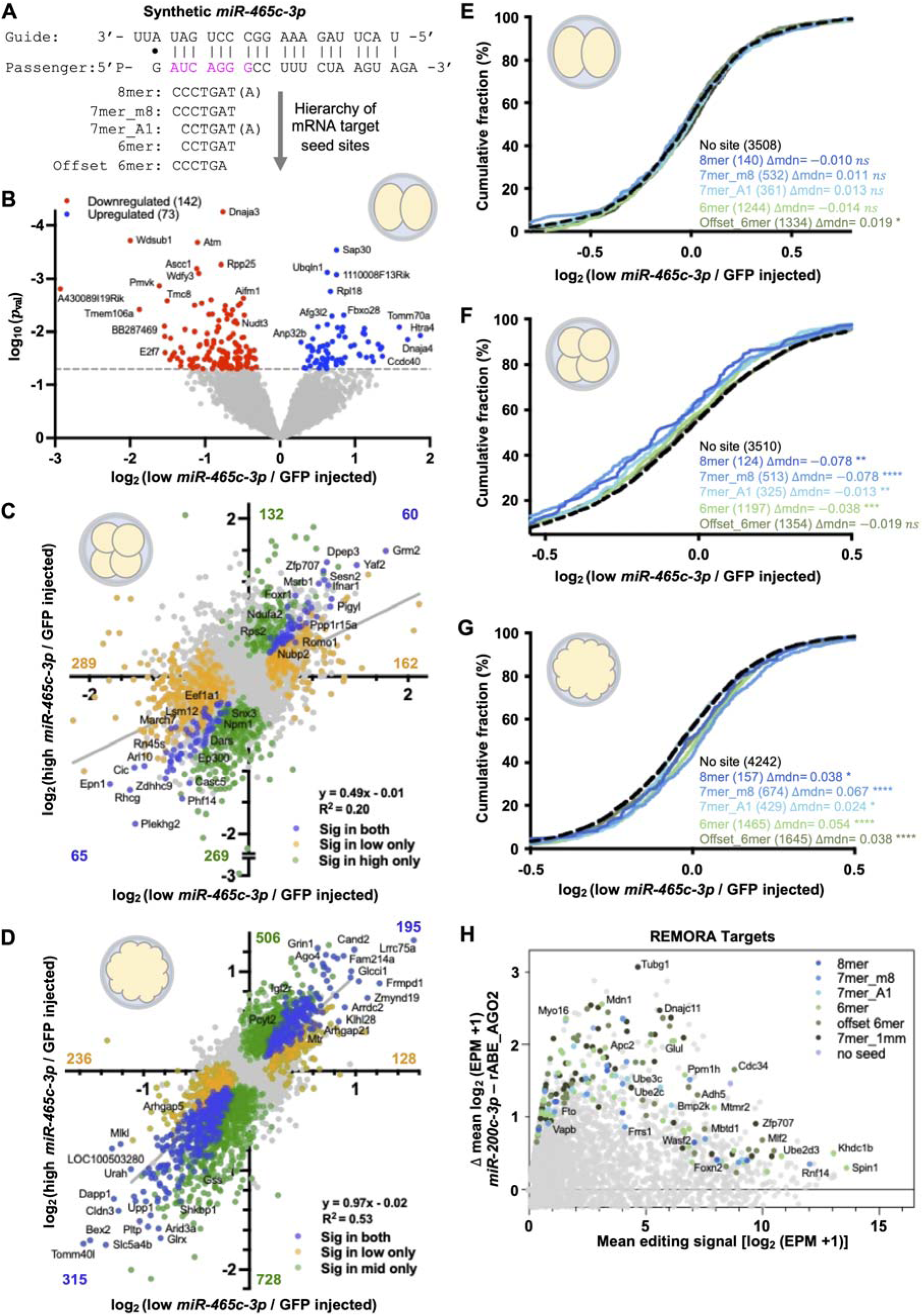
Different sperm miRNAs define distinct, seed-sequence–specific regulatory programs in early embryos. **(A)** Design of synthetic RNA engineered to preferentially load *miR-465c-3p*. **(B)** Volcano plot of DESeq2-derived differential gene expression in parthenogenetic embryos at the 2-cell stage following low-dose *miR-465c-3p* injection relative to GFP–injected controls. **(C-D)** Correlation of log_2_ fold change (DESeq2) between low-dose (x-axis) and high-dose (y-axis) *miR-465c-3p* injection relative to GFP–injected controls in 4-cell **(C)** and morula **(D)** stage parthenotes. Genes with nominal *p* < 0.05 are highlighted as indicated in legend. (**E-G)** ECDF of log_2_ fold changes (DESeq2) after low-dose *miR-465c-3p* injection for genes stratified by mutually exclusive *miR-465c-3p* seed-site class in 3’ UTRs in 2-cell **(E)**, 4-cell **(F)**, and morula **(G)** stage parthenotes. Distributional shifts relative to genes lacking canonical seed sites were tested using Wilcoxon rank-sum tests. Median differences relative to the no- canonical-seed baseline are reported. Asterisks denote enrichment significance *(*p* < 0.05), **(*p* < 0.01), ***(*p* < 0.001), ****(*p* < 0.0001). **(H)** MA plot showing gene level editing abundance in low-dose *miR-465c-3p*–injected 2-cell parthenotes. The x-axis shows the average log_2_(EPM+1) across rABE_AGO2 and rABE_AGO2+*miR-465c-3p* parthenotes, and the y-axis represents the difference in average log2(EPM+1) between conditions (rABE_AGO2+*miR-465c-3p* minus rABE_AGO2). Genes exhibiting significantly increased editing upon *miR-465c-3p* addition were identified using a one-sided Wilcoxon rank-sum test on embryo-level log_2_(EPM+1) values and are highlighted as indicated in the legend. Highlighted genes have nominal *p* < 0.05.

Unexpectedly, cumulative distribution analysis revealed that *miR-465c-3p* does not result in seed-dependent repression at the 2-cell stage (Figure 6E). In contrast, low-dose injection produced pronounced downregulation of seed-containing transcripts at the 4-cell stage (Figure 6F) which was exacerbated with high-dose injection (Supplemental Figure 4A). Additionally, genes with noncanonical seed sites were also globally downregulated by low-dose injection at the 4-cell stage (Supplemental Figure 4B). At the morula stage, compensatory upregulation of genes with seed sites was identified (Figure 6F and Supplemental Figure 4C), indicating that dose-sensitive target engagement is induced later in embryonic development for *miR-465c-3p* than what was observed for *miR-200c-3p*.

Despite *miR-465-3p* low-dose injection not inducing detectable direct downregulation of seeded genes at the 2-cell stage, AGO2-REMORA profiling revealed 167 of 171 transcripts exhibiting *miR-465c-3p*–dependent increases in RNA editing above baseline AGO2 activity contain either canonical or noncanonical seed sites, consistent with sequence-dependent direct recruitment of AGO2 to endogenous target RNAs (Figure 6G, Supplemental 4D). However, amongst these targeted genes, there was no detectable downregulation at the 2-cell or 4-cell stage with the low-dose injection (Supplemental Figure 4E-F). While we suspect that an increased dose of *miR-465c-*3p injection may induce detectable downregulation of targets even at the 2-cell stage, another alternative explanation is *miR-465c-3p* may be regulating these genes via translational regulation^32,33^. Nonetheless, AGO2-REMORA identified targets have reliably increased editing induced by injection of *miR-465c-3p* across detectable editing ranges (Supplemental Figure 4G).

Like *miR-200c-3p,* many AGO2-REMORA defined direct targets of *miR-465-3p* encode regulatory proteins, including 13 transcription factors, 12 RNA-binding proteins, and 8 chromatin-associated factors. These regulatory factors, overlap very little with the genes regulated by *miR-200c-3p* (Supplemental Figure 3E). Thus, sperm miRNAs target unique cohorts of regulatory factors which can establish secondary gene expression regulation during subsequent development. Together, these results demonstrate that dose-dependent, seed-sequence-defined miRNA targeting and repression represent a generalizable regulatory principle in early embryos, extending beyond *miR-200c-3p*. These comparative analyses reveal that distinct sperm miRNAs encode unique, dose-dependent regulatory trajectories, underscoring the specificity with which individual small RNAs can influence embryonic gene regulatory networks.

## DISCUSSION

These findings demonstrate for the first time that the early embryo is acutely sensitive to sperm-derived miRNA input, with measurable consequences for gene regulation that extend through preimplantation development and culminate in organismal phenotypes. Importantly, sperm miRNA abundance is not fixed; paternal environmental exposures, including diet, stress, and alcohol consumption, have been shown to alter the levels of specific sperm miRNAs. By titrating individual sperm miRNAs across physiologically relevant concentrations, we show that variation on the order of hundreds of molecules is sufficient to elicit reproducible, sequence-specific gene expression responses whose magnitude and scope scale with miRNA dose. Together, these results establish that sperm-delivered miRNAs function not as binary on/off switches, but as tunable regulatory inputs capable of quantitatively shaping developmental gene expression and transcriptional trajectories during the earliest stages of development. This quantitative sensitivity has clear biological relevance, given that sperm miRNA abundance varies in response to paternal environmental exposures, such that even modest differences in miRNA molecule number may constitute biologically meaningful signals transmitted to the embryo, linking paternal experience to altered in developmental outcomes.

Mechanistically, this exquisite sensitivity is mediated through canonical Argonaute engagement in the early embryo. Using AGO2–REMORA, we provide direct molecular evidence that sperm miRNAs recruit Argonaute to endogenous embryonic transcripts in a seed-sequence-dependent manner at the 2-cell stage. These early targeting events are linked to mRNA destabilization and precede stage-specific regulatory remodeling observed at later developmental transitions. Together, these data support a model in which transient miRNA activity at zygotic genome activation initiates regulatory cascades that are developmentally interpreted and progressively elaborated across preimplantation development (Figure 4E).

Our findings refine, rather than contradict, prior conclusions that miRNA activity is globally suppressed in oocytes and early embryos^34^. Early genetic ablation of *Dgcr8*, an essential miRNA biogenesis factor, demonstrated that canonical miRNAs are largely dispensable for oocyte maturation and cleavage-stage progression, leading to the prevailing view that miRNA-mediated regulation is functionally minimal during preimplantation development^34^. However, those studies were designed to detect overt developmental failure rather than subtle quantitative modulation of gene regulatory states. These observations led to the conventional view that miRNA-mediated regulation is either inactive or functionally irrelevant during early embryogenesis, and by extension that sperm-delivered miRNAs are unlikely to exert meaningful effects at fertilization. However, such studies were designed to detect overt developmental defects or widespread transcriptomic disruption, not subtle, quantitative regulatory effects whose consequences become apparent only over subsequent developmental stages. Our data instead support a model in which miRNA activity in the early embryo operates below the threshold required to disrupt core developmental programs yet remains sufficient to bias gene regulatory trajectories in ways that are progressively reinforced across development.

Importantly, comparison of distinct sperm miRNAs revealed that these responses are sequence-specific and temporally distinct. *miR-200c-3p* and *miR-465c-3p* each engaged endogenous targets in a seed-dependent manner yet exhibited divergent dose sensitivities and developmental trajectories. This specificity argues against a generic small RNA-mediated effect and instead supports a model in which individual sperm miRNAs encode discrete regulatory information that is interpreted by the embryo in a stage-dependent fashion.

Finally, we demonstrate that delivery of physiologically calibrated copy numbers of a single sperm miRNA, *miR-200c-3p*, is sufficient to induce reproducible craniofacial alterations in offspring. These phenotypes recapitulate features associated with paternal alcohol exposure in mice, directly linking early miRNA perturbation to later developmental outcomes. While additional environmental and genetic factors undoubtedly modulate these effects, our findings provide experimental support for the concept that environmentally sensitive changes in sperm miRNA content influence offspring phenotype.

Together, our results redefine the functional window of miRNA activity in the early embryo and resolve how small, stoichiometrically modest contributions from sperm can exert lasting developmental consequences. By acting as transient yet direct regulators at fertilization, sperm miRNAs encode structured information that initiates regulatory cascades that unfold across embryogenesis, thereby linking paternal experience to offspring phenotype.

## RESOURCE AVAILABILITY

### Lead Contact

Further information and requests for resources and reagents should be directed to and will be fulfilled by the lead contact, Colin Conine.

### Materials Availability

Plasmids generated in this study and additional information required to reproduce the experiments reported in this paper are available from the Lead Contact upon request.

### Data and code availability

The raw and processed sequencing data discussed in this publication have been deposited in NCBI Gene Expression Omnibus and are accessible via the accession GEO: GSE322558. Additionally, processed sequencing data is available as Supplemental Tables 1-6.

## ACKNOWLEDGEMENTS

We thank Yizhu Lin and Stephen Floor for generously sharing RNA editing enzyme constructs prior to publication of REMORA, enabling key aspects of this study. We are grateful to Marisa S. Bartolomei, Rebecca A. Simmons, and Taku Kambayashi for critical review of the manuscript and constructive feedback. We are also grateful to members of the Conine laboratory and Prithvi Sinha for helpful discussions and ongoing support. This work was supported by a Pew Biomedical Scholars Award to C.C.C., a Eunice Kennedy Shriver National Institute of Child Health and Human Development F31HD114433 and Contraception and Infertility Research L50HD119810 to G.S.L, an NSF GRFP to T.M.E, and NIH/NIAA R01AA028219 to M.C.G.

## AUTHOR CONTRIBUTIONS

G.S.L and C.C.C. conceived and designed the experiments. G.S.L. performed the experiments. G.S.L., J.G., and C.C.C analyzed data. S.L.H., K.Z.S., M.C.G., performed analysis of E16.5 embryos. A.H. performed surgical embryo transfers. G.S.L, T.M.E., M.L., C.C.C. collected E16.5 data. M.L. and L.S. managed mouse colonies. G.S.L. drafted the original manuscript with contributions from C.C.C., M.C.G., J.G. C.C.C. supervised the study.

## KEY RESOURCES TABLE

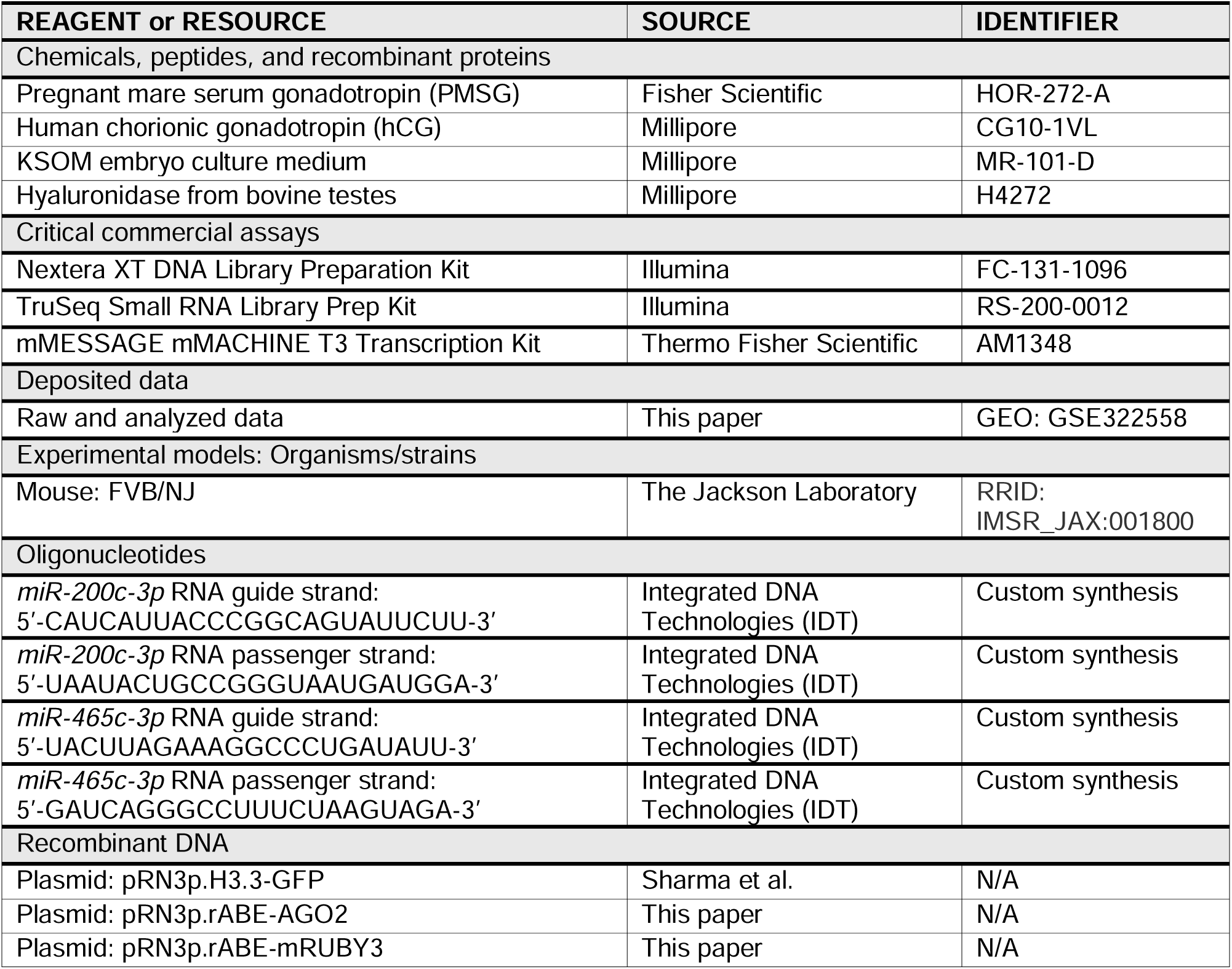
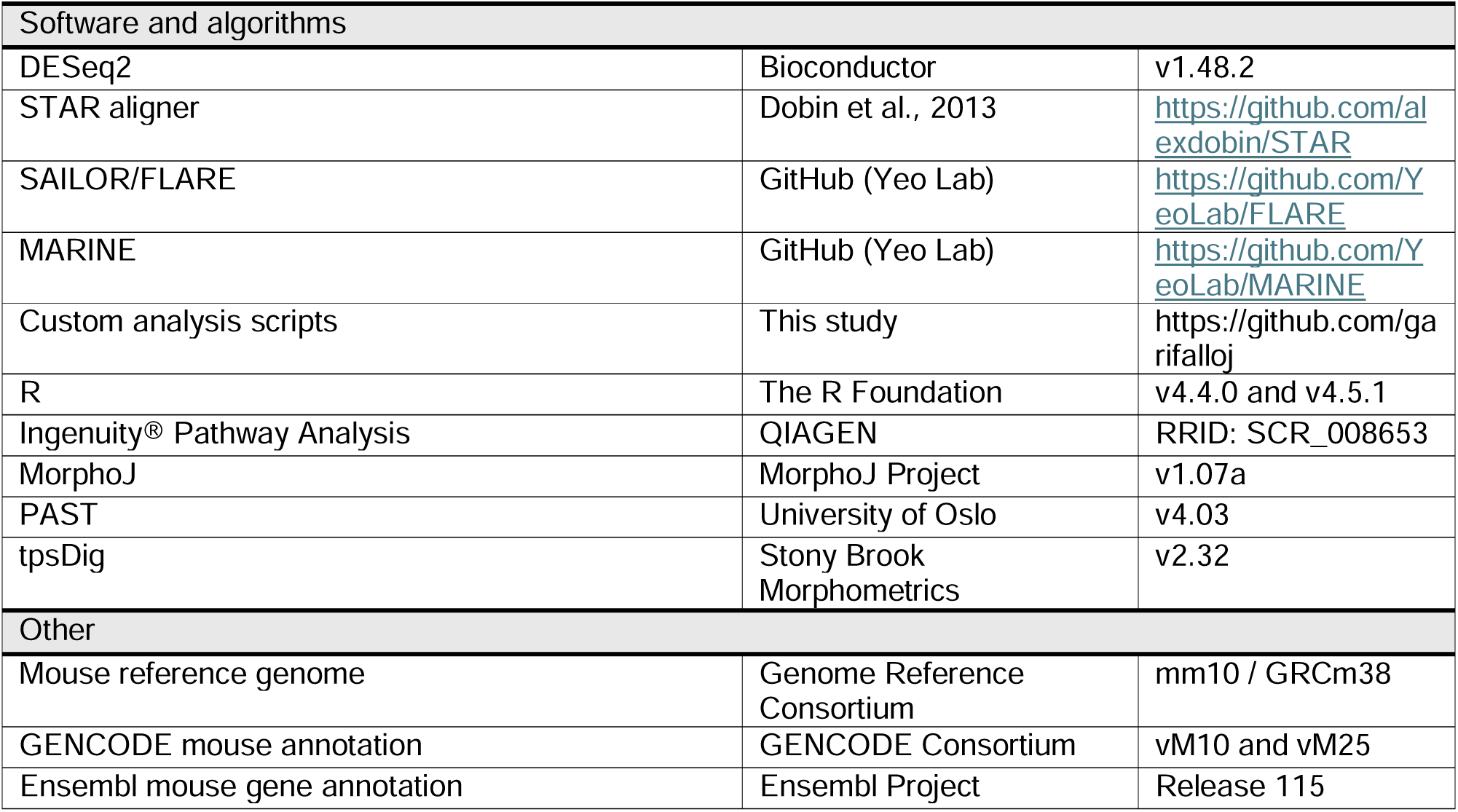

