## Supplemental Information for "Physiologic variation in sperm miRNAs tune embryonic gene regulatory programs and developmental outcomes"

### Supplemental Figures

#### Supplemental Figure 1

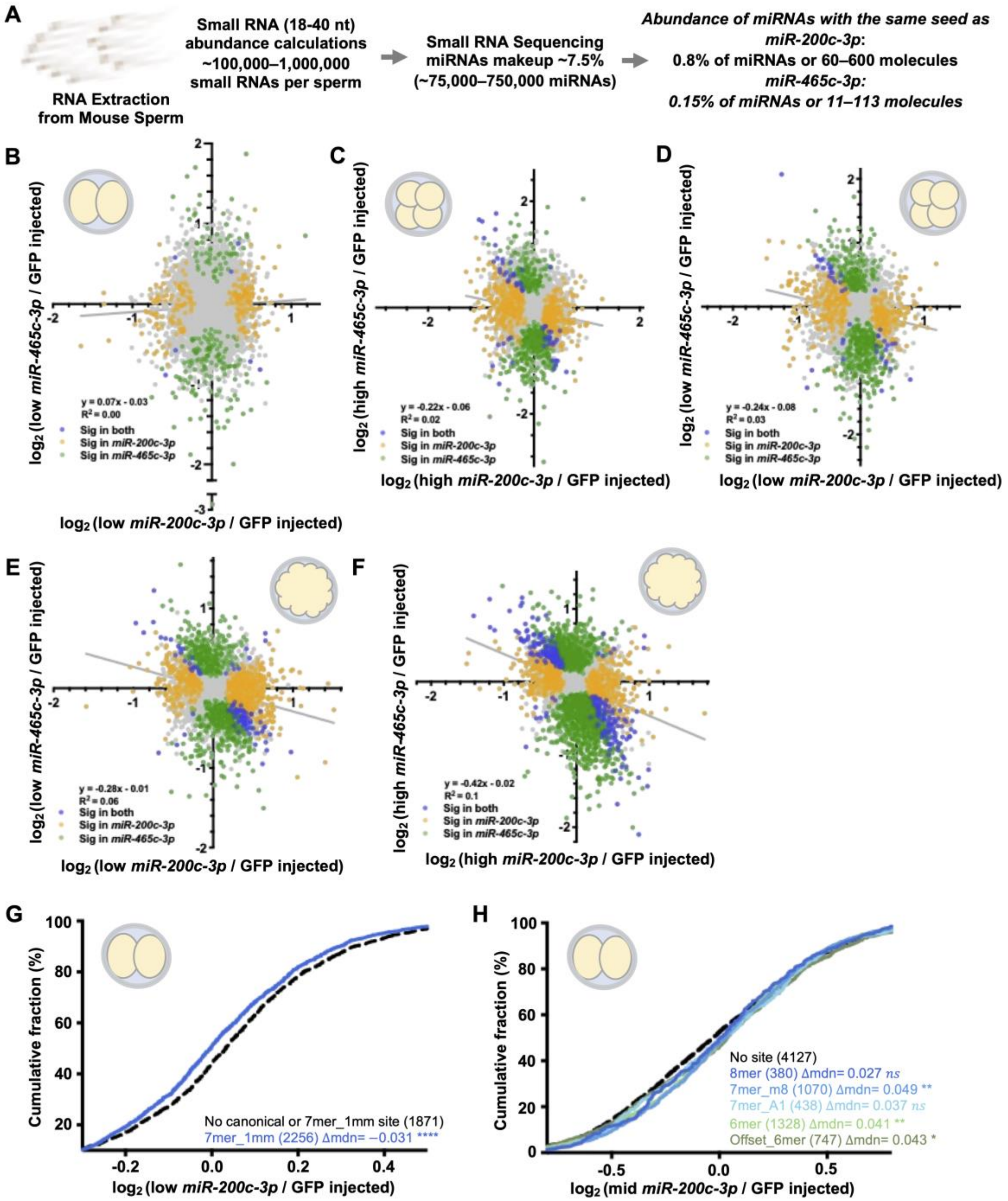

**Supplemental Figure 1. Distinct miRNAs drive unique dose-dependent, seed-sequence directed gene expression regulation in 2-cell parthenotes. (A)** Overview of how physiologic levels of sperm miRNAs were estimated. **(B-F)** Correlation of $\log_2$ fold change (DESeq2) between low-dose *miR-200c-3p* (x-axis) and low-dose (y-axis) *miR-465c-3p* injection relative to controls in 2-cell **(B)**, 4-cell **(D)**, and morula **(E)** parthenotes. Correlation between high-dose *miR-200c-3p* (x-axis) and high-dose (y-axis) *miR-465c-3p* injection relative to controls in 4-cell **(C)**, and morula **(F)** parthenotes. Genes with nominal $p < 0.05$ are highlighted as indicated in legend. **(G)** ECDF of $\log_2$ fold change (DESeq2) at the 2-cell stage after low-dose *miR-200c-3p* injection for genes with a 7mer_1mm seed-site in their 3' UTR. Distributional shifts relative to genes lacking canonical or 7mer_1mm seed sites were tested using Wilcoxon rank-sum tests. Median differences relative to no canonical or noncanonical seed baseline are reported. **(H)** ECDF of $\log_2$ fold change (DESeq2) at the 2-cell stage after mid-dose *miR-200c-3p* injection stratified by mutually exclusive *miR-200c-3p* seed-site class within 3' UTRs. Distributional shifts relative to genes lacking canonical seed were tested using Wilcoxon rank-sum tests. Median differences relative to no-canonical-seed baseline are reported. Asterisks denote significance *($p < 0.05$), **($p < 0.01$), ***($p < 0.001$), ****($p < 0.0001$).

### Supplemental Figure 2

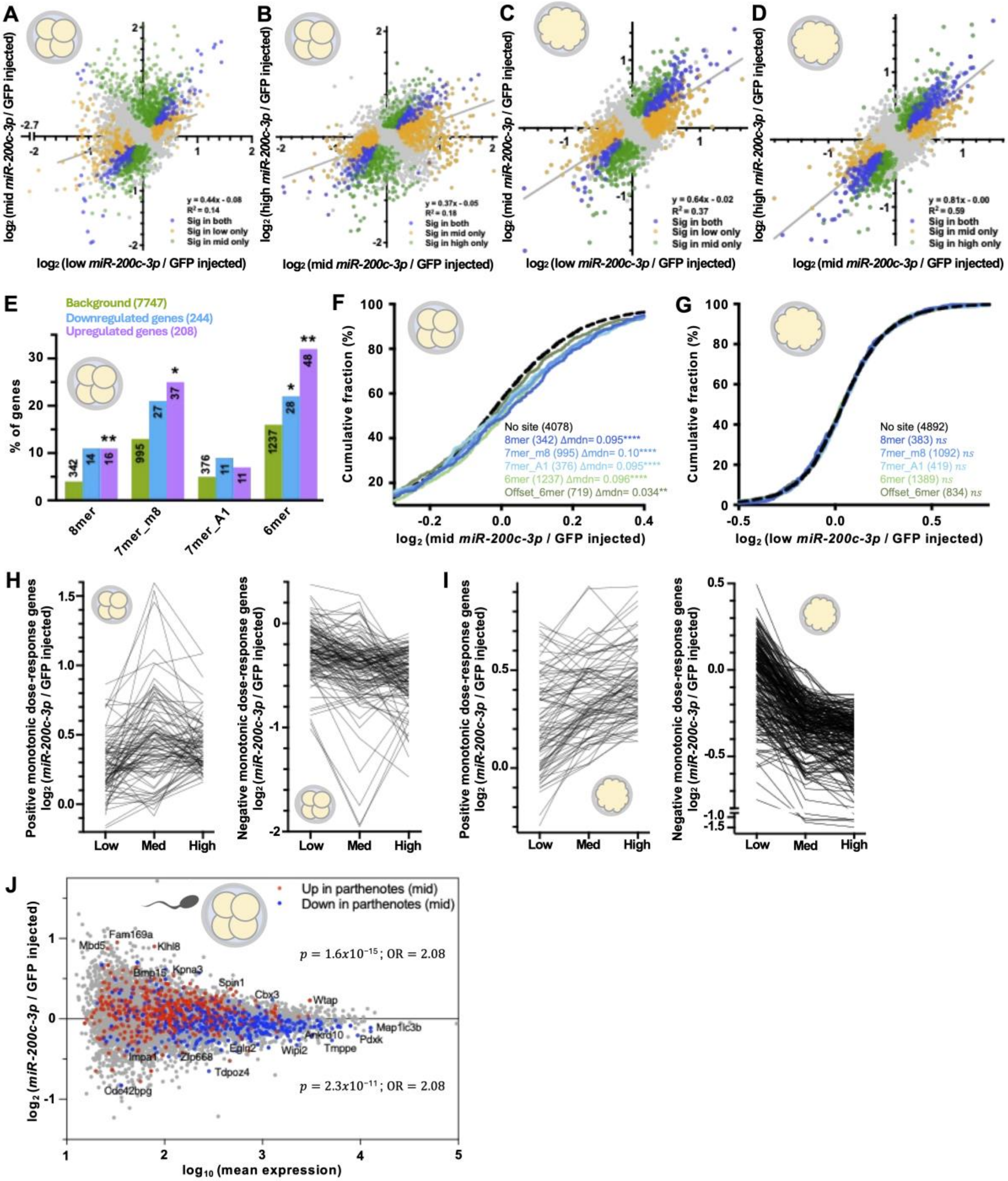

**Supplemental Figure 2. Tunable, dose-dependent transcriptional responses initiated by *miR-200c-3p* propagate across preimplantation development. (A-E)** Correlation of $\log_2$ fold change (DESeq2) between low (x-axis) and mid-dose (y-axis) *miR-200c-3p* injection relative to controls in 4-cell **(A)** and morula **(C)** parthenotes. Correlation of $\log_2$ fold change (DESeq2) between mid (x-axis) and high-dose (y-axis) *miR-200c-3p* injection relative to controls in 4-cell **(B)** and morula **(D)** parthenotes. **(E)** Proportion of genes that are upregulated and downregulated by *miR-200c-3p* low-dose injection containing *miR-200c-3p* seed sites in their 3' UTRs. Fisher's exact test $p-values$ are shown. **(F)** ECDF of $\log_2$ fold changes in 4-cell parthenotes for genes stratified by mutually exclusive *miR-200c-3p* seed site class in 3' UTRs following mid-dose injection. **(G)** ECDF of $\log_2$ fold changes in morula-stage parthenotes for genes stratified by mutually exclusive *miR-200c-3p* seed site class in 3' UTRs following low-dose injection. Distributional shifts relative to genes lacking canonical seed sites were tested using Wilcoxon rank-sum tests. Median differences relative to no-canonical-seed baseline are reported. **(H-I)** Distribution of $\log_2$ fold changes (DESeq2) for genes exhibiting positive (left) or negative (right) monotonic dose-response behavior across low-, mid-, and high-dose *miR-200c-3p* injection at the 4-cell **(H)** and morula **(I)** stages. Dose-dependent genes were identified using a likelihood ratio test (LRT; nominal $p < 0.05$), and the direction of dose response was classified using a numeric-dose Wald test. n = 88 activated and 124 repressed genes in 4-cells; n = 88 activated and 219 repressed genes in morula. **(J)** MA plot of gene expression changes at the 4-cell stage in *miR-200c-3p*–injected sperm-fertilized IVF embryos relative to GFP–injected controls. Genes that significantly dysregulated in 4-cell parthenogenetic embryos following mid-dose *miR-200c-3p* injection are highlighted. Fisher's exact test statistics, odds ratios (OR) and two-sided $p-values$, for comparisons between genes up- and down-regulated in parthenotes are shown. Asterisks denote significance *($p < 0.05$), **($p < 0.01$), ***($p < 0.001$), ****($p < 0.0001$).

### Supplemental Figure 3

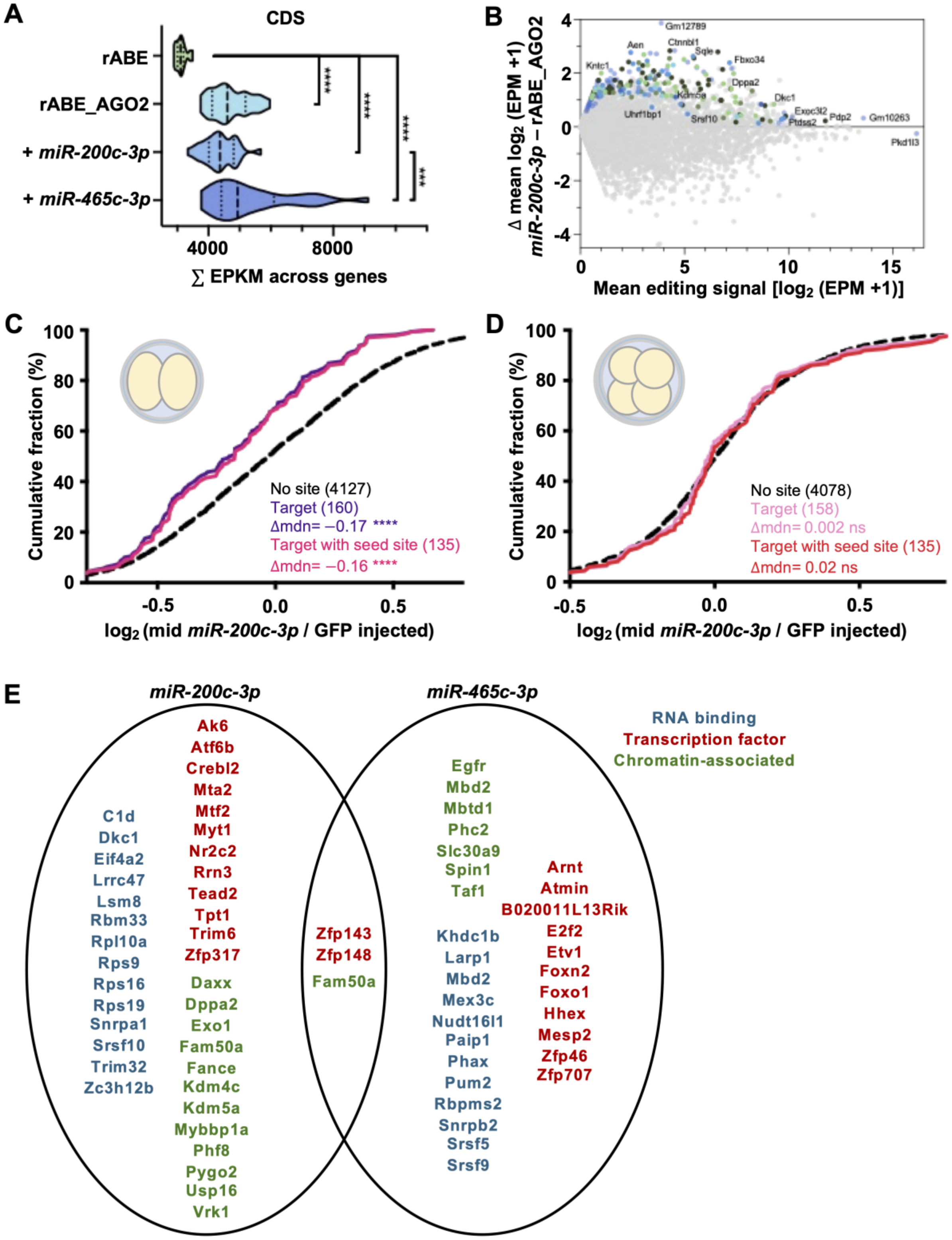

**Supplemental Figure 3. AGO2-REMORA identifies direct *miR-200c-3p* target mRNAs in early embryos** (**A)** Distribution of summed editing events per kilobase mapped reads (EPKM) per replicate across experimental conditions in mRNA coding sequences (CDS). **(B)** MA plot showing gene level editing abundance in low-dose 2-cell parthenotes. The x-axis shows the average $\log_2$(EPM+1) across rABE_AGO2 and rABE_AGO2+*miR-200c-3p* parthenotes, and the y-axis represents the difference in average log2(EPM+1) between conditions (rABE_AGO2+*miR-200c-3p* minus rABE_AGO2). Genes exhibiting significantly increased editing upon *miR-200c-3p* addition were identified using a one-sided Wilcoxon rank-sum test on embryo-level $\log_2$(EPM+1) values and are highlighted as indicated in the legend. Highlighted genes have nominal $p < 0.05$**.** **(C-D)** ECDF of $\log_2$ fold-changes (DESeq2) in 2-cell **(C)** and 4-cell **(D)** following mid-dose *miR-200c-3p* injection in parthenotes compared to control for target mRNAs identified by AGO2-REMORA. Distributional shifts relative to genes lacking canonical seed were tested using Wilcoxon rank-sum tests. Median differences relative to no-canonical-seed baseline are reported. Asterisks denote significance *($p < 0.05$), **($p < 0.01$), ***($p < 0.001$), ****($p < 0.0001$).

### Supplemental Figure 4

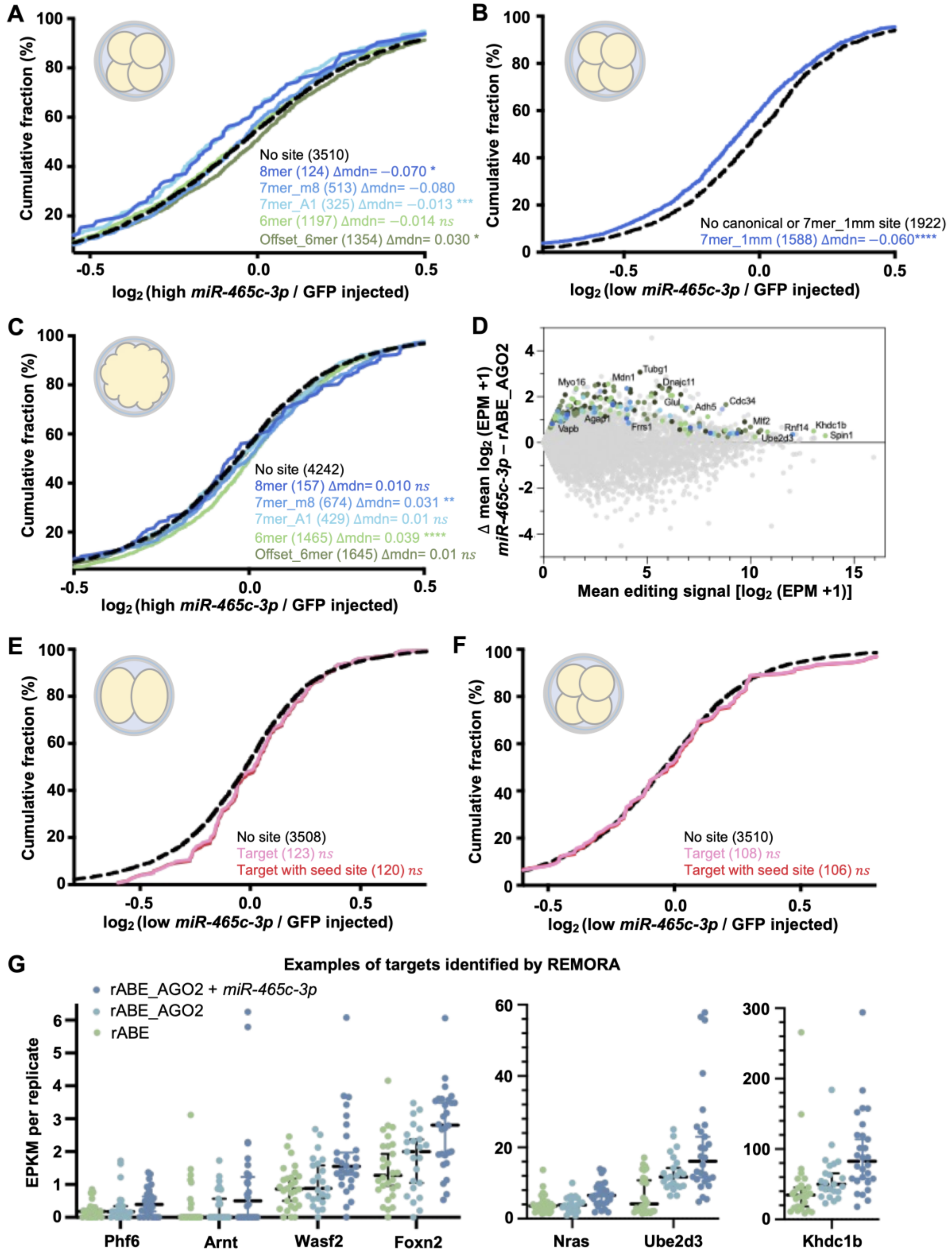

**Supplemental Figure 4. Different sperm miRNAs define distinct, seed-sequence–specific regulatory programs in early embryos (A,C)** ECDF of $\log_2$ fold-changes (DESeq2) from mRNA-seq data in 4-cell **(A)** and morula **(C)** following high-dose *miR-465c-3p* injection into parthenotes compared to controls, stratified by mutually exclusive *miR-465c-3p* seed-site class within 3' UTRs. **(B)** ECDF of $\log_2$ fold change (DESeq2) at the 4-cell stage after low-dose *miR-465c-3p* injection for genes with a 7mer_1mm seed-site in their 3' UTR. Distributional shifts relative to genes lacking canonical and/or noncanonical seed sites as indicated were tested using Wilcoxon rank-sum tests. Median differences relative to no-canonical-seed baseline are reported. Asterisks denote significance *($p < 0.05$), **($p < 0.01$), ***($p < 0.001$), ****($p < 0.0001$). **(D)** MA plot showing gene level editing abundance in low-dose 2-cell parthenotes. The x-axis shows the average $\log_2$(EPM+1) across rABE_AGO2 and rABE_AGO2+*miR-465c-3p* parthenotes, and the y-axis represents the difference in average log2(EPM+1) between conditions (rABE_AGO2+*miR-465c-3p* minus rABE_AGO2). Genes exhibiting significantly increased editing upon *miR-465c-3p* addition were identified using a one-sided Wilcoxon rank-sum test on embryo-level $\log_2$(EPM+1) values and are highlighted as indicated in the legend. Highlighted genes have nominal $p < 0.05$**.** **(E-F)** ECDF of $\log_2$ fold-changes (DESeq2) in 2-cell **(E)** and 4-cell **(F)** following low-dose *miR-465c-3p* injection in parthenotes compared to control for target mRNAs identified by AGO2-REMORA. Distributional shifts relative to genes lacking canonical seed were tested using Wilcoxon rank-sum tests. **(G)** Gene-level editing signal (EPKM) in 2-cell parthenotes for select target mRNAs identified by AGO2-REMORA containing canonical seed sequences. Median with 95% confidence intervals is indicated. Y-axis scales differ between panels.

### Supplemental Data

**Supplemental Data 1. Procrustes ANOVA for craniofacial shape differences in the frontal profile of E16.5 male and female offspring derived from embryos injected with *miR-200c-3p* compared with controls.** Sample sizes: male *miR-200c-3p* (n=15), male control (n=16), female *miR-200c-3p* (n=9), female control (n=11)

| **Centroid size:** | | | | | |
|---|---|---|---|---|---|
| Effect | SS | MS | df | F | P (param.) |
| Individual | 90.04538 | 30.015513 | 3 | 0.65 | 0.5898 |
| Residual | 2186.204542 | 46.514990 | 47 | | |
| **Shape, Procrustes ANOVA:** | | | | | |
| Effect | SS | MS | df | F | P (param.) |
| Individual | 0.06723936 | 0.0002490347 | 270 | 1.20 | 0.0167 |
| Residual | 0.87853079 | 0.0002076905 | 4230 | | |

**Supplemental Data 2. Procrustes ANOVA for craniofacial shape differences in the left profile of E16.5 male and female offspring derived from embryos injected with *miR-200c-3p* compared with controls.** Sample sizes: male *miR-200c-3p* (n=14), male control (n=16), female *miR-200c-3p* (n=10), female control (n=11)

| **Centroid size:** | | | | | |
|---|---|---|---|---|---|
| Effect | SS | MS | df | F | P (param.) |
| Individual | 113.891940 | 37.963980 | 3 | 0.51 | 0.6799 |
| Residual | 3524.783162 | 74.995386 | 47 | | |
| **Shape, Procrustes ANOVA:** | | | | | |
| Effect | SS | MS | df | F | P (param.) |
| Individual | 0.27661417 | 0.0002490347 | 270 | 1.20 | 0.0167 |
| Residual | 0.76148614 | 0.0001653248 | 4606 | | |

**Supplemental Data 3. Procrustes ANOVA for craniofacial shape differences in the right profile of E16.5 male and female offspring derived from embryos injected with *miR-200c-3p* compared with controls.** Sample sizes: male *miR-200c-3p* (n=15), male control (n=15), female *miR-200c-3p* (n=10), female control (n=10)

| **Centroid size:** | | | | | |
|---|---|---|---|---|---|
| Effect | SS | MS | df | F | P (param.) |
| Individual | 218.501702 | 72.833901 | 3 | 0.84 | 0.4782 |
| Residual | 3981.449947 | 86.553260 | 46 | | |
| **Shape, Procrustes ANOVA:** | | | | | |
| Effect | SS | MS | df | F | P (param.) |
| Individual | 0.06637516 | 0.0002257659 | 294 | 1.41 | <0.0001 |
| Residual | 0.72321029 | 0.0001604282 | 4508 | | |

**Supplemental Data 4. Procrustes ANOVA for craniofacial shape differences in the frontal profile of E16.5 male offspring derived from embryos injected with *miR-200c-3p* compared with controls.** Sample sizes: *miR-200c-3p* (n=15), control (n=16).

| **Centroid size:** | | | | | |
|---|---|---|---|---|---|
| Effect | SS | MS | df | F | P (param.) |
| Individual | 8.806632 | 8.806632 | 1 | 0.16 | 0.6949 |
| Residual | 1627.402672 | 56.117334 | 29 | | |
| **Shape, Procrustes ANOVA:** | | | | | |
| Effect | SS | MS | df | F | P (param.) |
| Individual | 0.02954095 | 0.0003282327 | 90 | 1.53 | 0.0012 |
| Residual | 0.56143127 | 0.0002151078 | 2610 | | |

**Supplemental Data 5. Procrustes ANOVA for craniofacial shape differences in the left profile of E16.5 male offspring derived from embryos injected with *miR-200c-3p* compared with controls.** Sample sizes: *miR-200c-3p* (n=14), control (n=16).

| **Centroid size:** | | | | | |
|---|---|---|---|---|---|
| Effect | SS | MS | df | F | P (param.) |
| Individual | 5.583089 | 5.583089 | 1 | 0.06 | 0.8095 |
| Residual | 2640.020931 | 94.286462 | 28 | | |

| Shape, Procrustes ANOVA: | | | | | |
|---|---|---|---|---|---|
| Effect | SS | MS | df | F | P (param.) |
| Individual | 0.05896743 | 0.0006017085 | 98 | 3.05 | <0.0001 |
| Residual | 0.54149067 | 0.0001973363 | 2744 | | |

**Supplemental Data 6. Procrustes ANOVA for craniofacial shape differences in the right profile of E16.5 male offspring derived from embryos injected with *miR-200c-3p* compared with controls.** Sample sizes: *miR-200c-3p* (n=15), control (n=15).

| Centroid size: | | | | | |
|---|---|---|---|---|---|
| Effect | SS | MS | df | F | P (param.) |
| Individual | 33.051798 | 33.051798 | 1 | 0.30 | 0.5876 |
| Residual | 3075.171767 | 109.82763 | 28 | | |
| **Shape, Procrustes ANOVA:** | | | | | |
| Effect | SS | MS | df | F | P (param.) |
| Individual | 0.00896311 | 0.0000914603 | 98 | 0.52 | 1 |
| Residual | 0.48028184 | 0.0001750298 | 2744 | | |

**Supplemental Data 7. Procrustes ANOVA for craniofacial shape differences in the frontal profile of E16.5 female offspring derived from embryos injected with *miR-200c-3p* compared with controls.** Sample sizes: *miR-200c-3p* (n=9), control (n=11).

| Centroid size: | | | | | |
|---|---|---|---|---|---|
| Effect | SS | MS | df | F | P (param.) |
| Individual | 0.000069 | 0.000069 | 1 | 0.00 | 0.9988 |
| Residual | 558.801870 | 31.044548 | 18 | | |
| **Shape, Procrustes ANOVA:** | | | | | |
| Effect | SS | MS | df | F | P (param.) |
| Individual | 0.01364485 | 0.0001516094 | 90 | 0.77 | 0.9420 |
| Residual | 0.31772924 | 0.0001961292 | 1620 | | |

**Supplemental Data 8. Procrustes ANOVA for craniofacial shape differences in the left profile of E16.5 female offspring derived from embryos injected with *miR-200c-3p* compared with controls.** Sample sizes: *miR-200c-3p* (n=10), control (n=11).

| Centroid size: | | | | | |
|---|---|---|---|---|---|
| Effect | SS | MS | df | F | P (param.) |
| Individual | 0.004845 | 0.004845 | 1 | 0.00 | 0.9920 |
| Residual | 884.762231 | 46.566433 | 19 | | |
| **Shape, Procrustes ANOVA:** | | | | | |
| Effect | SS | MS | df | F | P (param.) |
| Individual | 0.02572862 | 0.0002625370 | 98 | 2.26 | <0.0001 |
| Residual | 0.21617301 | 0.0001160972 | 1862 | | |

**Supplemental Data 9. Procrustes ANOVA for craniofacial shape differences in the right profile of E16.5 female offspring derived from embryos injected with *miR-200c-3p* compared with controls.** Sample sizes: *miR-200c-3p* (n=10), control (n=10).

| **Centroid size:** | | | | | |
|---|---|---|---|---|---|
| Effect | SS | MS | df | F | P (param.) |
| Individual | 0.192815 | 0.192815 | 1 | 0.00 | 0.9513 |
| Residual | 906.278181 | 50.348788 | 18 | | |
| **Shape, Procrustes ANOVA:** | | | | | |
| Effect | SS | MS | df | F | P (param.) |
| Individual | 0.01411024 | 0.0001439821 | 98 | 1.04 | 0.3701 |
| Residual | 0.24353308 | 0.0001380573 | 1764 | | |
